# Internalisation of integrin-bound extracellular matrix modulates invasive carcinoma cell migration

**DOI:** 10.1101/2024.01.11.575153

**Authors:** Montserrat Llanses Martinez, Keqian Nan, Zhe Bao, Rachele Bacchetti, Shengnan Yuan, Joe Tyler, Xavier Le Guezennec, Frédéric A. Bard, Elena Rainero

**Affiliations:** School of Biosciences, University of Sheffield, Western Bank, Sheffield, S10 2TN, UK; Institute of Molecular and Cell Biology, 61 Biopolis Drive, Proteos, Singapore 138673, Singapore; Centre de Recherche en Cancérologie de Marseille, CRCM, Marseille, France

## Abstract

The interaction between cancer cells and the extracellular matrix (ECM) plays a pivotal role in tumour progression. While the extracellular degradation of ECM proteins has been well characterised, ECM endocytosis and its impact on cancer cell progression, migration and metastasis is poorly understood. ECM internalisation is increased in invasive breast cancer cells, suggesting it may support invasiveness. Here we developed a high-content screening assay to study ECM uptake. We identified that mitogen-activated protein kinase (MAPK) family members, MAP3K1 and MAPK11 (p38β), and the protein phosphatase 2 (PP2) subunit PPP2R1A were required for the internalisation of ECM-bound α2β1 integrin. Furthermore, α2β1 integrin was necessary for macropinocytosis of soluble dextran, identifying it as a novel and targetable regulator of macropinocytosis in cancer. Moreover, disruption of α2 integrin, MAP3K1, MAPK11 and PP2R1A-mediated ECM internalisation significantly impaired cancer cell migration and invasion in 2D and 3D culture systems. Finally, α2β1 integrin and MAP3K1 expression were significantly upregulated in pancreatic tumours and correlated with poor prognosis in pancreatic cancer patients. Strikingly, MAP3K1, MAPK11, PPP2R1A and α2 integrin expression were higher in chemotherapy-resistant tumours in breast cancer patients. Our results identified the α2β1 integrin/p38 signalling axis as a novel regulator of ECM endocytosis, which drives invasive migration and tumour progression.

---

The extracellular matrix (ECM) is a non-cellular 3D structure that surrounds cells and organs *in vivo*. Depending on its organisation, it is classified into basement membrane and interstitial matrix. The ECM dynamics are tightly regulated in morphogenesis, differentiation and tissue homeostasis; therefore, dysregulations in ECM remodelling are associated with pathological conditions, including fibrosis and tumour progression. Cells engage with the ECM through the heterodimeric integrin receptors, which are composed of an α and a β subunit and activate diverse pro-survival signal transduction pathways^1,2^. However, the ECM may simultaneously pose a physical constraint for cancer cell invasion and tumour initiation^3,4^. As a result, ECM degradation accompanies tumour progression. Our lab has recently shown that ECM internalisation supports proliferation of breast cancer cells under amino acid starvation by promoting tyrosine metabolism^5^, through a mechanism that requires ECM macropinocytosis, followed by lysosomal degradation^5^. However, the signalling regulators controlling ECM endocytosis are poorly understood.

α2β1 integrin is the major collagen I receptor and is expressed in epithelial cells, platelets and fibroblasts, among other cell types^6^. α2β1 integrin mediates the reorganisation and contraction of collagen matrices in fibroblasts^7^. In addition, α2β1 integrin has been shown to regulate melanoma and pancreatic carcinoma cell migration^8,9^, as well as bone metastatic dissemination of prostate and breast cancer cells^10,11^. However, the mechanisms underlying α2β1 integrin-mediated cancer cell invasive migration have not been elucidated. Integrin/ECM binding triggers the activation of several signalling pathways, including the mitogen-activated protein kinase (MAPK) pathway. In mammalian cells, there are four distinct MAPK signalling pathways according to the terminal tier kinase: extracellular signal-regulated kinase (ERK) 1/2, c-Jun N-terminal Kinase (JNK) 1/2/3, p38 and ERK5^12,13^. p38 MAPK is a family of four serine/threonine-specific protein kinases: p38α or MAPK14, p38β or MAPK11, p38γ or MAPK12, and p38δ or MAPK13^14^. p38α and p38β are ubiquitously expressed and share 75% amino acid sequence homology^14^. p38 MAPK plays an important role in cell proliferation, differentiation, stress responses, autophagy and cell migration, among other biological processes^14^.

A deep comprehension of how cancer cells interact with the ECM is essential to understand tumour growth and metastasis. Here we showed that ECM internalisation was upregulated in primary cancer cells extracted from a polyoma middle T (PyMT)-driven mouse model of breast cancer, compared to non-transformed mammary epithelial cells and ECM components were trafficked through EEA1-positive early endosomes and delivered to the lysosomes. Through a kinase and phosphatase functional screen, we identified MAP3K1, MAPK11 and PPP2R1A as novel regulators of the macropinocytosis of ECM components, which we defined as being mediated by α2β1 integrin. Interestingly, invasive breast cancer cells were shown to internalise ECM as they migrated on cell-derived matrices (CDMs) and blocking ECM uptake impaired the invasive migration of breast, pancreatic and ovarian cancer cells, both in 2D and 3D culture systems. Finally, α2β1 integrin and MAP3K1 were significantly upregulated in pancreatic tumours compared to healthy tissue and high expression of these genes correlated with reduced survival of pancreatic cancer patients. Remarkably, chemotherapy-resistant breast tumours showed higher mRNA expression levels of MAP3K1, MAPK11, PPP2R1A and α2 integrin. Altogether we identified a novel signalling pathway linking ECM macropinocytosis, degradation and invasive migration in different cancer types, which could potentially be exploited for the generation of novel therapeutic interventions to prevent metastatic dissemination.

## Results

### A kinase and phosphatase screen identifies MAPK signalling cascade as a novel regulator of Matrigel internalisation in breast cancer cells

Polyoma middle T oncogene expression under the mammary epithelial MMTV promoter constitutes a widely used mouse model of breast cancer that recapitulates many aspects of the human disease^15,16^. To assess whether ECM endocytosis is upregulated in primary breast cancer cells, NMuMG cells, derived from normal mouse mammary glands, and PyMT#1 cells, generated from MMTV-PyMT tumours^15^, were seeded on fluorescently labelled Matrigel, a basement-membrane formulation, and CDM, generated by telomerase-immortalised human fibroblasts. It is widely established that, following endocytosis, the ECM is degraded in the lysosomes^17^. Therefore, to prevent lysosomal degradation and discern between changes that could be due to altered ECM degradation rather than endocytosis, cells were treated with a cysteine-cathepsin lysosomal inhibitor, E64d^5,18^. Internalisation of Matrigel was upregulated in PyMT#1 cells compared to NMuMG cells, both in the presence and the absence of E64d (**Extended data Fig. 1a**). Similarly, CDM internalisation was significantly higher in PyMT#1 cells in the presence of E64d in agreement with previous results^5^ (**Extended data Fig. 1b**). Conversely, E64d did not increase internalisation of CDM in NMuMG, suggesting that CDM lysosomal degradation is specifically triggered in invasive breast cancer cells (**Extended data Fig. 1b**). These data indicate that ECM internalisation and degradation is upregulated in invasive breast cancer.

**Fig. 1.**
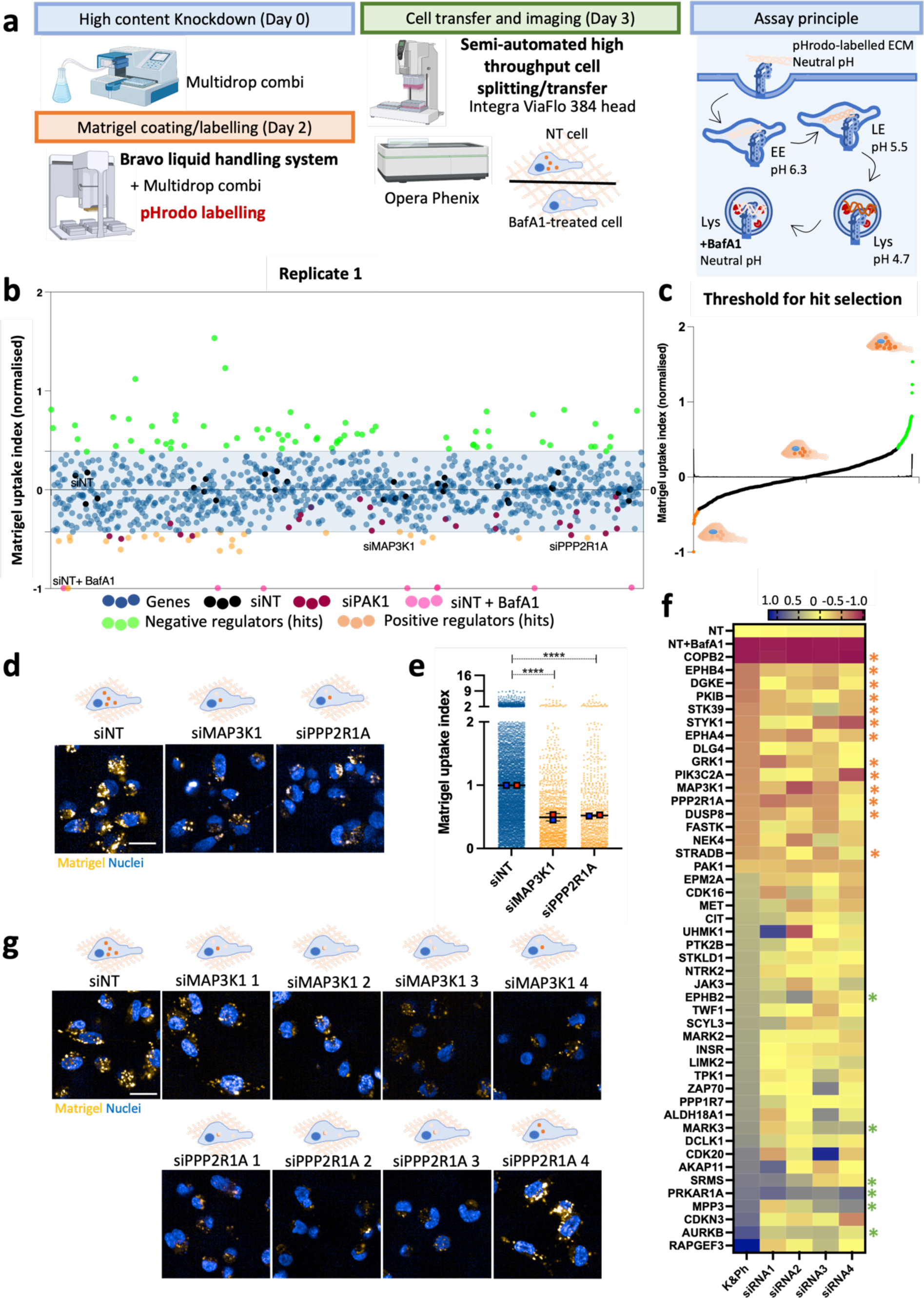
A kinome and phosphatome screen identified MAP3K1 and PPP2R1A as positive regulators of matrigel internalisation in MDA-MB-231 cells. **a,** Screen schematic representation. 3000 MDA-MB-231 cells were transfected, transferred into pH-rodo labelled 0.5mg/ml matrigel, incubated for 6h and labelled with lμg/ml Hoechst for nuclear staining. Cell imaging was carried out with an Opera phenix microscope (40X objective). Columbus software was used for image analysis, **b,** Normalised cloud plot analysis from replicate 1, positive regulator hits are in orange, negative hits in green, **c,** First derivative of the curve for hit threshold for replicate 1. **d,** Representative images from positive regulators in the screen, MAP3K1 and PPP2R1A. Scale bar, 20μm. **e,** Normalised cell data quantification for MAP3K1 and PPP2R1A. Data are presented as the normalised mean ± SD; N=3 independent replicates. ****p<0.0001; Kruskal-Wallis test, **f,** Heatmap of kinome and phosphatome siRNA deconvolution. N=2 technical replicates, N=2 biological replicates per individual siRNA. Orange stars identified validated positive regulators, green stars validated negative regulators, g, Representative images of the deconvoluted siRNA for MAP3K1 and PPP2R1A.

To determine the trafficking pathway followed by ECM components, we measured Matrigel colocalization with an early endosomal marker, EEA1, and a late endosomal/lysosomal marker, LAMP2, in MDA-MB-231 triple-negative breast cancer cells. Matrigel colocalisation with EEA1 progressively decreased over time, with the strongest colocalization observed 3h after seeding the cells, as indicated by the colocalization map (**Extended data Fig. 1c**). In contrast, Matrigel colocalisation with LAMP2 increased over time and reached a strong colocalisation after 12h (**Extended data Fig. 1d**). These data indicate that Matrigel is first delivered to early endosomes and subsequently to lysosomes for degradation. The endosomal-lysosomal system creates an enclosed environment which is progressively acidified as cargos are transported towards the lysosomes^19^. This difference in pH can be exploited to specifically visualise cargos within the endosomal system, by using pH-sensitive dyes. MDA-MB-231 cells were seeded on collagen I labelled with NHS-fluorescein and a pH-sensitive dye, pHrodo, which fluorescence increases as the endosomal compartment is acidified^20,21^. Using time-lapse microscopy, we showed that as collagen I is internalised, there is a progressive increase in the intensity of pHrodo, while the NHS-fluorescein signal is slightly decreased, consistent with the quenching of the green fluorescence in acidic environments^22^ (**Supplementary video 1** and **Extended data Fig. 1e**). These data indicated that following endocytosis, endosomes are rapidly acidified (in about 12 minutes) and thus pHrodo is a reliable dye for dissecting ECM uptake mechanisms.

To identify the signalling pathways responsible for controlling ECM uptake, we screened an siRNA library targeting 948 protein kinases and phosphatases^23^. MDA-MB-231 cells were knocked down for 72h, transferred and seeded on pHrodo-labelled Matrigel for 6h, stained with a nuclear dye and imaged live (**Fig. 1a**). ECM uptake was normalised between the non-targeting (NT) and NT in presence of bafilomycin A1, a V-ATPase inhibitor that prevents lysosomal acidification, as a positive control (**Fig. 1b,c, Extended data Fig. 2a,b** and **Supplementary Table 1**). The Z-factor (Z’) provides insight into the quality of a high-throughput assay (where 1.0 > Z ≥ 0.5 indicates an excellent assay and 0.5 > Z > 0 is considered a marginal assay)^24^. The Z’ robust and Z’ standard calculated were respectively 0.6 and 0.694, indicating the high quality of the assay (**Extended data Fig. 2f**). Moreover, we observed a good correlation between replicates, with R^2^=0.6219 (**Extended data Fig. 2c**). Since we previously showed that p21-activated kinase 1 (PAK1) was required for ECM macropinocytosis^5^, PAK1 was included as an additional positive control. It was reassuring that PAK1 knock-down consistently reduced Matrigel uptake in the screen (**Extended data Fig. 2d,e**).

**Fig. 2.**
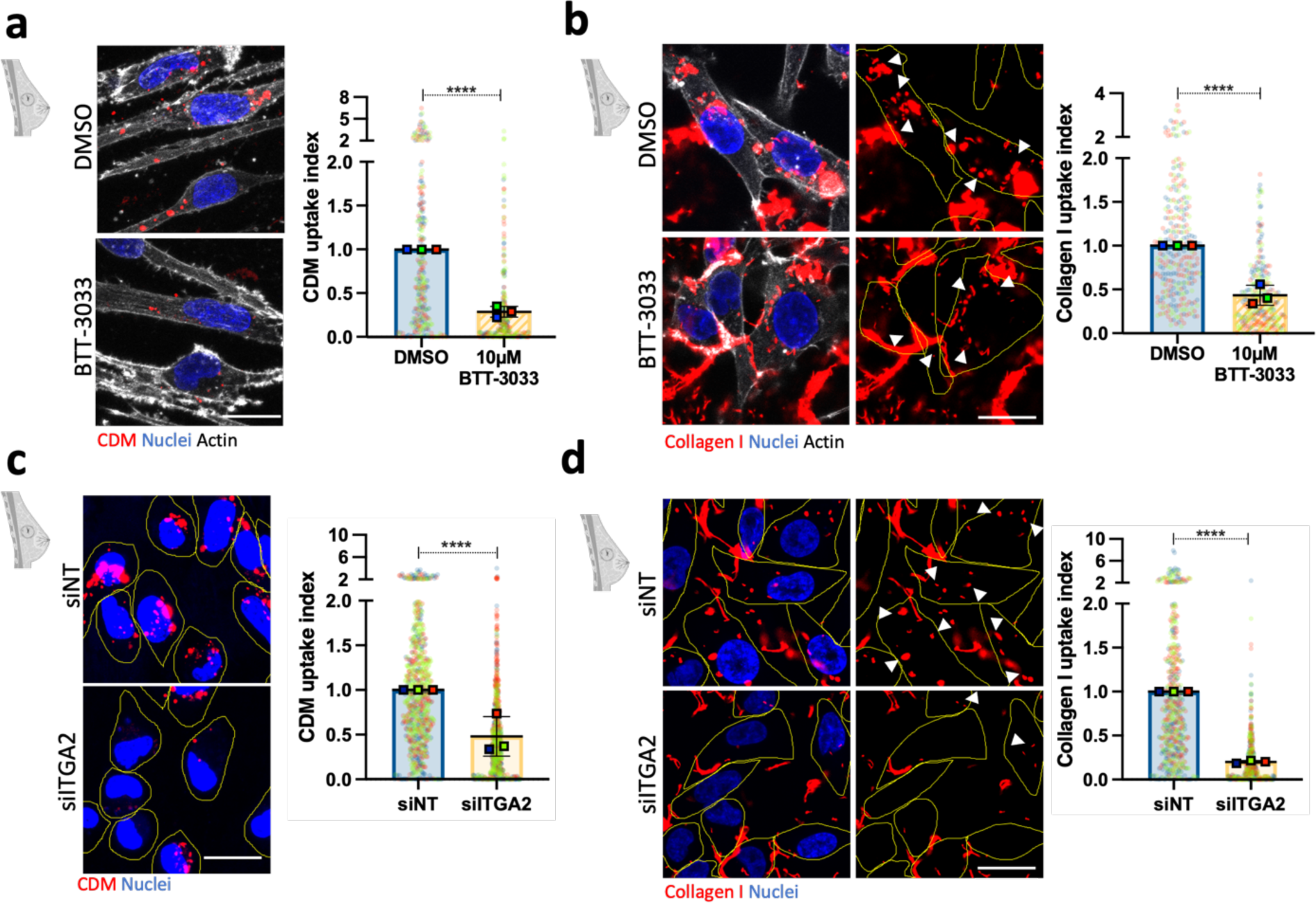
α2 integrin promoted ECM internalisation in MDA-MB-231 breast cancer cells. **a,** MDA-MB-231 cells were allowed to adhere on biotinylated CDM for 2h before incubating the cells with 10μM BTT-3033, an α2 integrin inhibitor, or DMSO for 6h. Cells were fixed and stained with phalloidin Alexa-Fluor 555, Alexa fluor 488-streptavidin and DAPI. Data are presented as the normalised mean ± SD; N=3 independent replicates. ****p<0.0001; Mann-Whitney test, **b,** MDA-MB-231 cells were cultured on NHS Alexa Fluor 555-labelled 1mg/ml collagen I for 2h before treatment with 10μM BTT-3033, or DMSO for an additional period of 6h. Cells were fixed and stained for actin and nuclei. Data are presented as the normalised mean ± SD; N=3 independent replicates. ****p<0.0001; Mann-Whitney test, **c,** MDA-MB-231 cells were transfected with an siRNA targeting α2 integrin (silTGA2) or a non-targeting siRNA control (siNT), seeded on pHrodo-labelled CDM for 6h, stained with Ipg/ml Hoechst and imaged live. Data are presented as the normalised mean ± SD; N=3 independent replicates. ****p<0.0001; Mann-Whitney test, **d,** MDA-MB-231 cells transfected as in **c,** seeded on Alexa Fluor 555-labelled 1mg/ml collagen I for 6h, fixed and stained for nuclei. Data are presented as the normalised mean + SD; N=3 independent replicates. ****p<0.0001; Mann-Whitney test.

We utilised Reactome^25^ to assess the enrichment for biological pathways in the positive and negative regulators identified by the screen, and enriched pathways included negative regulation of MAPK signalling pathway (p-value: 2.68E-05) and MAPK signalling cascades (p-value: 3.23E-04; **Supplementary Table 2**). MAP3K1 and PPP2R1A were top positive regulators in the screen (**Fig. 1d,e**). COPB2 knockdown, despite resulting in a strong inhibition in ECM uptake, was not considered further as it was associated with high levels of cell toxicity. MAP3K1 promotes ERK1/2, JNK and p38 activation under distinct stimuli^26,27^, while protein phosphatase 2 scaffold subunit α (PPP2R1A) is the regulatory subunit of protein phosphatase 2 A (PP2A)^28^. Consistent with the pathway analysis, we explored additional siRNA targets relevant for MAPK pathway and checked Matrigel uptake upon ERK1/2 (MAPK3 and MAPK1), JNK1/2/3 (MAPK8-10) and p38 (MAPK 11-14) knockdown in the screen. Only MAPK11 (p38β) consistently reduced Matrigel internalisation (**Extended data Fig. 2g,h**), suggesting that MAPK11 signalling is important in this process.

To validate our hits, we performed a secondary screen for 45 common hits obtained between replicates. The pooled siRNAs for each gene were deconvoluted into four individual siRNAs (**Supplementary Table 3**). We considered it an on target hit if 2 out of 4 siRNAs reduced or increased Matrigel uptake. Based on siPAK1, a 30% reduction or increase compared to NT was used as a cut-off to confirm positive or negative regulators. According to this criterion, 19 genes were validated, among them MAP3K1 and PPP2R1A (**Fig. 1f,g**). As in the primary screen, knocking down COPB2 strongly reduced both Matrigel uptake index and cell count, indicating cell toxicity. While the individual siRNAs against the positive regulators gave consistent results, this was not the case for most of the negative regulators (on-target hits marked by a green star in **Fig. 1f**). Therefore, we focused on the positive regulators of ECM uptake, including MAP3K1, MAPK11 and PPP2R1A.

### ECM internalisation is dependent on α2β1 integrin

Our screen identified MAPK signalling as an important regulator of ECM internalisation. Collagens have been linked to the activation of p38 MAPK in platelets^29^, while the collagen receptor α2β1 integrin activates p38 MAPK and PP2A in fibroblasts^30,31^. We therefore hypothesised that α2β1 integrin may activate MAPK11 and regulate ECM internalisation. Indeed, downregulation of β1 integrin significantly reduced Matrigel internalisation in MDA-MB-231 cells (**Extended data Fig. 3a,h**). To assess whether the ECM was trafficked together with β1 integrin, we assessed β1 integrin localisation in cells seeded on fluorescently labelled Matrigel. Colocalization analysis revealed a strong overlap between Matrigel and β1 integrin at all the time points measured (**Extended data Fig. 3b**). This data supports the hypothesis that β1 integrin not only regulates ECM endocytosis but is trafficked together with the ECM.

**Fig. 3.**
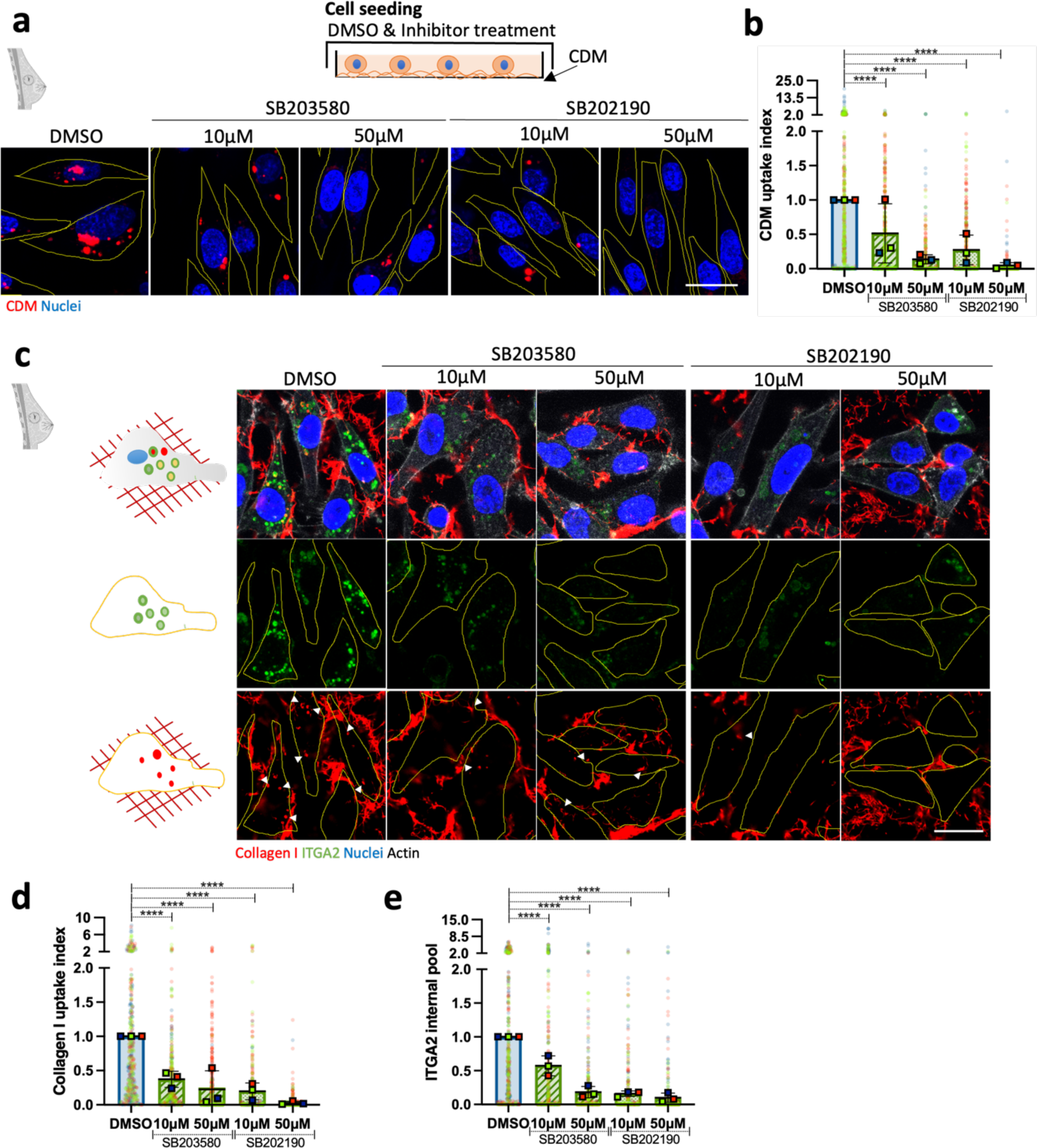
p38 MAPK regulated the internalisation of collagen l-occupied α2β1 integrin. **a,** MDA-MB-231 cells were serum starved for 16 to 18h. 3×10^5^ cells were seeded on pHrodo-labelled CDM for 6hr in the presence of DMSO, 10μM or 50μM of SB203580 and SB202190 in 5% FBS and imaged live. Scale bar, 22μm. **b,** CDM uptake index was calculated with ImageJ. Values represented are normalised mean + SD from N=3 experiment; ****p<0.0001; Kruskal-Wallis test, **c,** MDA-MB-231 cells were serum starved for 16 to 18h. 3×10^5^ cells were seeded on 1mg/ml collagen I, labelled with NHS-Alexa fluor 555, treated as in a, fixed and stained for α2 integrin (ITGA2) and nuclei. Scale bar, 20μm. **d,e** Collagen I uptake index and α2 integrin internal pool were calculated with ImageJ. Values represented are normalised mean + SD from N=3 independent experiments; ****p<0.0001; Kruskal-Wallis test.

In MDA-MB-231 cells, treatment with the α2 integrin pharmacological inhibitor BTT-3033, which disrupts collagen binding, significantly decreased CDM and collagen I endocytosis (**Fig. 2a,b**). Consistently, siRNA-mediated downregulation of α2 integrin reduced the internalisation of CDM and collagen I (**Fig. 2c,d**). We then wanted to determine whether α2-dependent ECM uptake was shared between different cancer types or was a specific feature associated with invasive breast cancer cells. In ovarian cancer, the overexpression of the small GTPase Rab25 promotes the internalisation of fibronectin-occupied α5β1 integrin to sustain invasive migration^32^; therefore we measured ECM internalisation in the highly invasive ovarian carcinoma cell line A2780, overexpressing Rab25 (A2780-Rab25). Similar to MDA-MB-231 cells, α2 integrin downregulation significantly reduced CDM and collagen I uptake in A2780-Rab25 cells (**Extended data Fig. 3c,d**). To study whether α2 integrin regulated this process in a primary cell line, we treated the polyoma middle T-driven mouse breast cancer cell line YEJ P^33^ with BTT-3033. Again, pharmacological inhibition of α2 integrin nearly abolished collagen I internalisation in YEJ P cells (**Extended data Fig. 3e**). Similar to breast tumours, pancreatic tumours are surrounded by an extremely dense and fibrotic stroma^34,35^ and pancreatic ductal adenocarcinoma (PDAC) cells have been previously shown to be able to internalise collagen I^36^. We assessed collagen I uptake in the metastatic PDAC cell line SW1990 and found that α2 integrin inhibition significantly reduced collagen I internalisation (**Extended data Fig. 3f**). Altogether, this data suggests that α2β1 integrin is required for ECM internalisation in different cancer types.

### MAP3K1, MAPK11 and PPP2R1A regulate macropinocytosis of ECM-bound α2β1 integrin

Our data suggested that MAPK signalling regulates ECM endocytosis (**Supplementary Table 2**). To characterise the role of MAPK signalling in this process, we screened four inhibitors against p38α/β (SB202190 and SB203580), ERK1/2 (FR180204) and MEK1/2 (PD98059) on Matrigel, collagen I and CDM uptake in MDA-MB-231 cells (**Extended data Fig. 4a**)^37–41^. The strongest uptake impairment was obtained by p38 inhibition in a dose-dependent manner, starting from 10µM (**Extended data Fig. 4b**). While 50µM ERK1/2 inhibition reduced ECM internalisation, inhibition of MEK1/2, an upstream activator of ERK1/2, did not reduce ECM uptake at effective inhibitory concentrations starting from 10µM (**Extended data Fig. 4b**). Altogether, this data supported a major role for p38 MAPK in mediating ECM internalisation.

**Fig. 4.**
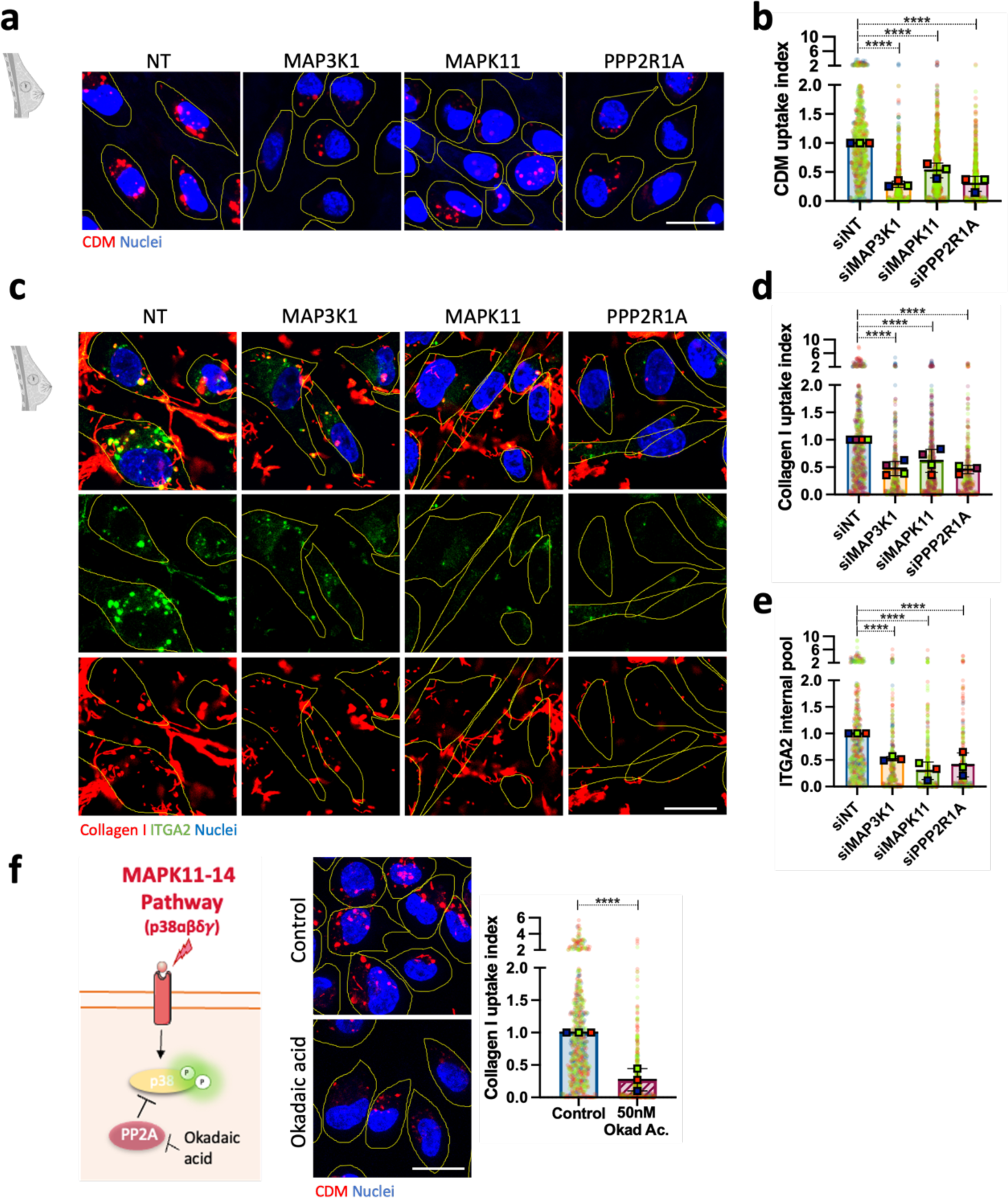
MAP3K1, MAPK11 and PPP2R1A modulated collagen I-occupied α2β1 integrin uptake. **a,** MDA-MB-231 cells were transfected with an siRNA targeting MAPK3Kl (siMAP3K1), an siRNA targeting MAPK11 (siMAPK11), an siRNA targeting PPP2R1A (siPPP2RlA) or a non-targeting siRNA control (siNT), seeded on pHrodo-labelled CDM for 6h, stained with 1µg/ml Hoechst and imaged live. Scale bar, 20µm. **b,** CDM uptake index was calculated with lmageJ. Values represented are normalised mean + SD from N=3 independent experiments; ****p<0.0001; Kruskal-Wallis test. **c,** MDA-MB-231 cells were transfected as in a, seeded on 1mg/ml collagen I, labelled with NHS-Alexa fluor 555, for 6h, fixed and stained for α2 integrin (ITGA2) and nuclei. Scale bar, 20µm. **d,e,** Collagen I uptake index and α2 integrin internal pool were calculated with lmageJ. Values represented are normalised mean+ SD from N=3 independent experiments; ****p<0.0001; Kruskal-Wallis test. **f,** 3×10^5^ MDA-MB-231 cells were cultured on pHrodo-labelled collagen I for 6hr in the presence of 50nM okadaic acid (Okad Ac.) or water (Control), stained with 1µg/ml Hoechst and imaged live. Scale bar, 20µm. Collagen I uptake index was measured with Image **J.** Values represented are normalised mean + SD from N=3 independent experiments; ****p<0.0001; Kruskal-Wallis test.

To corroborate these results, highly invasive MDA-MB-231 and A2780-Rab25 cells were seeded on labelled CDM and were treated with 10µM and 50µM of p38α/β inhibitors. Similarly, p38 inhibition reduced CDM internalisation in a dose-dependent fashion (**Fig. 3a,b; Extended data Fig. 4c**). These experiments were performed with pHrodo-labelled ECM. Under these conditions, a reduction in pHrodo signal could represent either a blockade of endocytosis, a defect in lysosomal targeting of internalised ECM or an impairment of lysosomal acidification. To address this, MDA-MB-231 cells and A2780-Rab25 cells were seeded on collagen I labelled with NHS-Alexa Fluor 555, a non-pH sensitive dye. Consistently, collagen I internalisation was markedly decreased upon p38α/β inhibition (**Fig.3c,d; Extended data Fig. 4d**), indicating that the observed effect was due to changes in endocytosis and not in endosomal acidification. Similarly, p38 inhibition reduced collagen I internalisation in YEJP and SW1990 cells (**Extended data Fig. 4e,f**), suggesting that p38 signalling regulates ECM internalisation in breast, ovarian and PDAC cells. Importantly, these tumours are characterised by a dense and collagen I-rich stroma. Since we showed that β1 integrin trafficked together with Matrigel and α2β1 integrin was required for ECM uptake (**Extended data Fig. 3b**), we aimed to assess whether p38 inhibition controlled the internalisation of α2β1 integrin, by measuring α2 internal pool. Indeed, p38 inhibition reduced the levels of internalised α2 integrin in MDA-MB-231 cells (**Fig.3c,e**). In agreement with our previous observations, α2 integrin could be detected in collagen I-positive vesicles (**Fig.3c**), indicating that p38 promotes the internalisation of ECM-bound α2β1 integrin.

To validate the positive regulators identified in our screen, we assessed CDM and collagen I uptake by highly invasive MDA-MB-231 and A2780-Rab25 cells in the presence of siRNA against MAP3K1, MAPK11 and PPP2R1A (**Extended data Fig. 5e-i**) and found that the knockdowns significantly reduced CDM and collagen I uptake in both cell lines (**Fig. 4a-d; Extended data Fig. 5a-d**). In MDA-MB-231 cells, downregulating MAP3K1, MAPK11 and PPP2R1A also significantly reduced α2 integrin internal pool (**Fig. 4c,e**). To rule out the possibility that the decrease in internalised α2 integrin was due to changes in its expression, we assessed how MAPK11 and PPP2R1A knockdown affected α2 integrin protein levels by western blotting. Knocking down PPP2R1A resulted in a ∼50% decrease in α2 integrin expression (**Extended data Fig. 5j**); while MAPK11 downregulation did not significantly affect α2 integrin levels (**Extended data Fig. 5k**). These data suggest that changes in the α2 integrin internal pool are likely due to reduced endocytosis upon MAPK11 downregulation; however, PPP2R1A may potentially regulate both α2 integrin trafficking and expression.

**Fig. 5.**
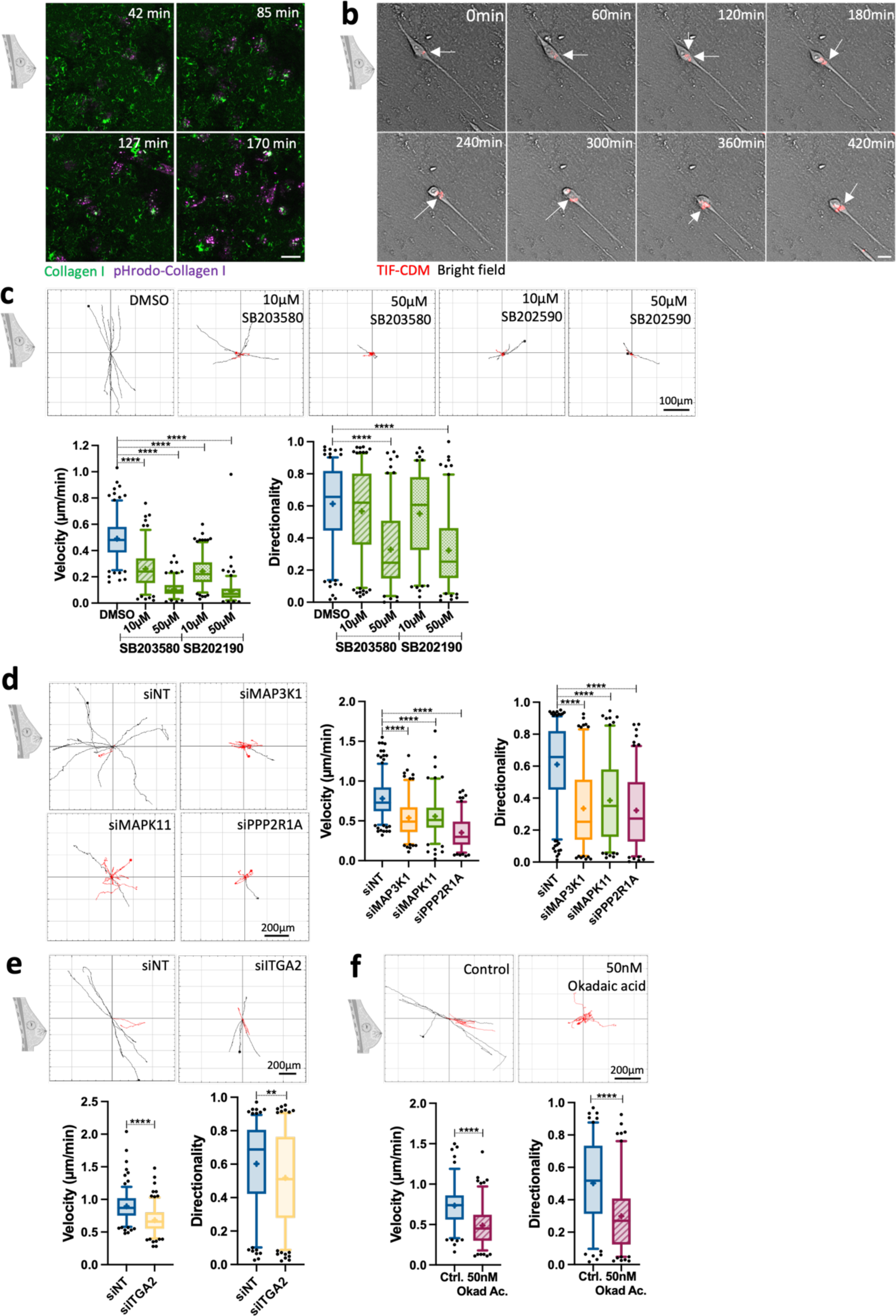
Regulators of ECM internalisation were required for invasive breast cancer cell migration. **a,** MDA-MB-231 cells were seeded on 1mg/ml collagen I labelled with pHrodo and NHS-fluorescein for 30 min before live imaging for 5h. Representative images extracted from Supplementary video 2 are shown. Scale bar, 20μm. **b,** MDA-MB-231 cells were seeded in pHrodo-labelled CDM for 6h and imaged live by time-lapse microscopy. Representative images extracted from supplementary video 3 are shown. Scale bar, 20μm. **c,** MDA-MB-231 cells were seeded on CDM for 6hr in the presence of DMSO, 10µM and 50µM SB203580 or SB202190 and imaged live with a 10X Nikon Inverted Ti eclipse with Oko-lab environmental control chamber for at least 7h. Spider plots show the migration paths of manually tracked cells (directionality >0.5 in black, <0.5 in red). Box and whisker plots represent 5-95 percentile, + represents the mean, dots are *<5%* and >95%; N= 3 independent experiments. ****p <0.0001; Kruskal Wallis test, **d,** MDA-MB-231 cells were transfected with an siRNA targeting MAP3K1 (siMAP3K1), an siRNA targeting MAPK11 (siMAPK11), an siRNA targeting PPP2R1A (siPPP2R1A) or a non-targeting siRNA control (siNT), seeded on CDM for 6hr and imaged live with a 10X Nikon Inverted Ti eclipse with Oko-lab environmental control chamber for 17h. Spider plots show the migration paths of manually tracked cells (directionality >0.5 in black, <0.5 in red). Box and whisker plots represent 5-95 percentile, + represents the mean; dots are <5% and >95%; N= 3 independent experiments. ****p <0.0001; Kruskal Wallis test, **e,** MDA-MB-231 cells were transfected with an siRNA targeting α2 integrin (silTGA2) or a non-targeting siRNA control (siNT), seeded on CDM for 4h and imaged live for 17h. Spider plots show the migration paths of manually tracked cells (directionality >0.5 in black, <0.5 in red). Box and whisker plots represent 5-95 percentile, + represents the mean, dots are <5% and >95%; N= 3 independent experiments. **p=0.0032; ****p<0.0001; Mann-Whitney test, f, MDA-MB-231 cells seeded on CDM for 6h in the presence of 50nM Okadaic acid (Okad Ac.) or vehicle control (water) and imaged live for 17h. Spider plots show the migration paths of manually tracked cells (directionality >0.5 in black, <0.5 in red). Box and whisker plots represent 5-95 percentile, + represents the mean, dots are <5% and >95%; N= 3 independent experiments. ****p<0.0001; Mann-Whitney test.

We have previously shown that breast cancer cells uptake different ECM components through mactopinocytosis^5^. To assess whether p38 was controlling this endocytic process, we measured dextran uptake in MDA-MB-231 cells seeded on collagen I (**Extended data Fig 6a**). Interestingly, p38 inhibition significantly reduced dextran internalisation (**Extended data Fig. 6b**), suggesting that this pathway is a conserved macropinocytic programme. Since integrins cluster in macropinocytic cups^42^, we reasoned α2 integrin may be required for macropinocytosis. Indeed, α2 inhibition with either BTT-3033 or siRNA-mediated downregulation significantly reduced dextran uptake (**Extended data Fig. 6c,d**), indicating that α2β1 integrin is a novel macropinocytosis regulator in invasive breast cancer cells.

**Fig. 6.**
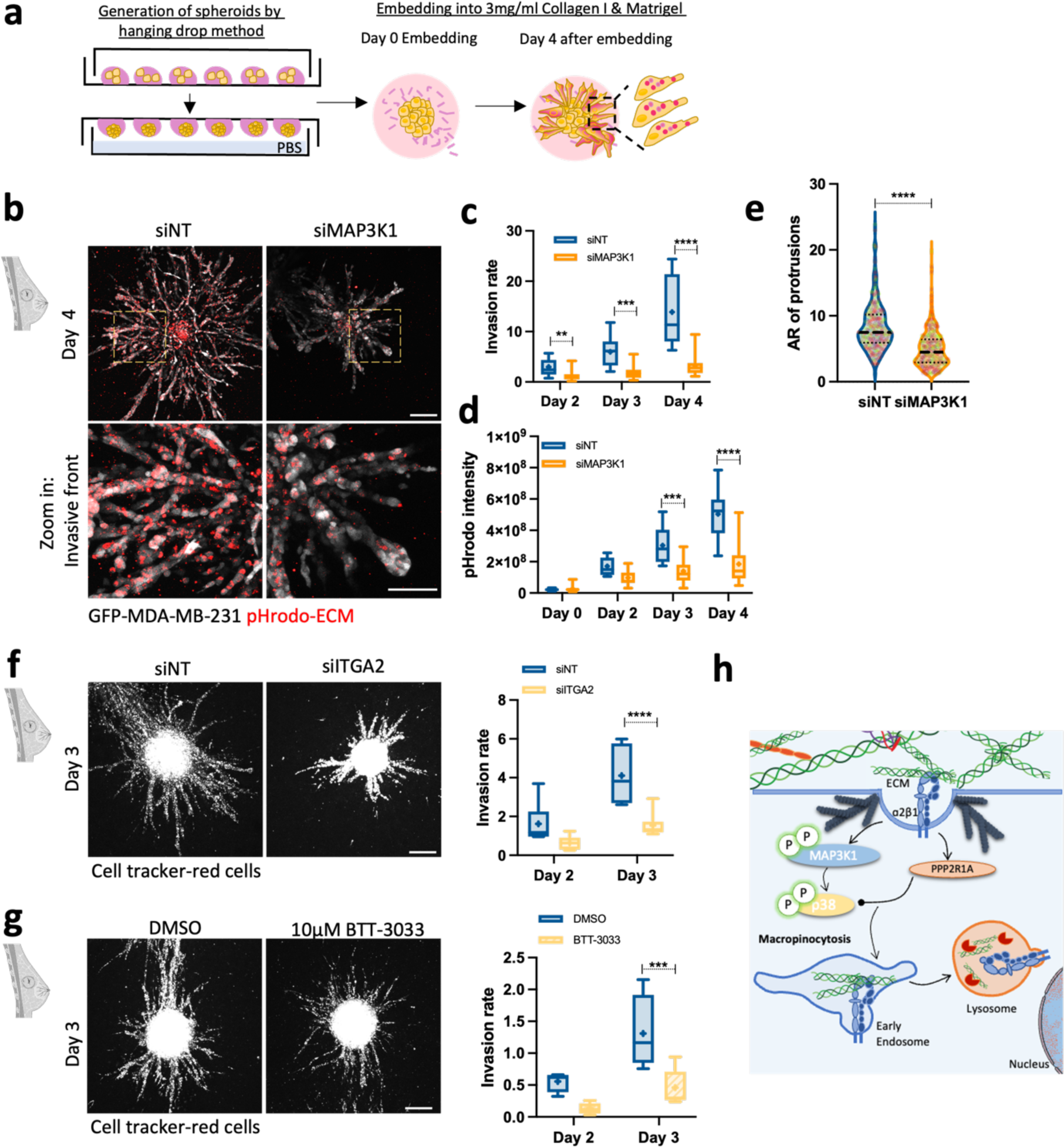
MAP3Kl and α2 integrin were necessary for MDA-MB-231 cell invasion. **a,** Schematic representation of ECM internalisation in 3D. **b,** MDA-MB-231 cells were transfected with an siRNA targeting MAP3Kl (siMAP3K1) or a non-targeting siRNA control (siNT) and spheroids embedded in 3D 3mg/ml pHrodo-labelled collagen I and matrigel (1:1 ratio) mixture for 4 days. Scale bar, 200µm. For the invasive front: scale bar, 100µm. **c,** Invasion rate (spheroid invasion area/spheroid core area). Box and whisker plots represent 5-95 percentile,+ represents the mean, dots are <5% and >95%; N= 4 independent experiments. **p=0.0060, ***p=0.0003, ****p <0.0001; 2-way ANOVA, Mixed-effects analysis test. d, pHrodo intensity in spheroids. Box and whisker plots represent Min to Max, + represents the mean; N= 4 independent experiments. ***p=0.0004, ****p <0.0001; Mixed-effects analysis test. **e,** Aspect ratio (AR) of invasive protrusions. Cell data are presented in violin plots as the median and quartiles; N=4 independent experiments. ****p <0.0001; Mann-Whitney test. **f,** MDA-MB-231 cells were transfected with an siRNA targeting a2 integrin (silTGA2) or non-targeting siRNA control (siNT) and spheroids were embedded in 2mg/ml collagen I and matrigel (1:1 ratio) matrix for 3 days. Scale bar, 200µm. Box and whisker plots represent Min to Max,+ represents the mean; N=2 independent experiments. ****p <0.0001; 2-way ANOVA test. **g,** MDA-MB-231 cells spheroids were embedded in 2mg/ml collagen I and matrigel (1:1 ratio) matrix and treated with DMSO or 10µM BTT-3033. Scale bar, 200µm. Box and whisker plots represent Min to Max,+ represents the mean; N=2 independent experiments. ***p=0.0005; 2-way ANOVA test.

Actin polymerisation by the WAVE complex drives macropinocytosis^42^ and PPP2R1A was described to form part of the WAVE shell complex, regulating actin polymerisation in a phosphatase activity-independent manner^43^. Simultaneously, PPP2R1A is the scaffold subunit of the protein phosphatase PP2A^44^. To discern whether PPP2R1A regulated ECM macropinocytosis through its phosphatase activity or via the regulation of actin polymerisation, we assessed collagen I uptake in the presence of okadaic acid, a PP2A inhibitor^45,46^. 50nM okadaic acid significantly reduced collagen I internalisation (**Fig. 4f**), suggesting that the phosphatase function of PPP2R1A regulated ECM uptake. Altogether, these data indicate that MAP3K1, MAPK11 and PPP2R1A control ECM uptake by modulating integrin endocytosis.

### Regulators of ECM internalisation modulate invasive cancer cell migration

Integrin endocytosis is essential for integrin turnover, migration and invasion^47^. Internalisation of fibronectin-bound α5β1 integrin is required for invasive migration in A2780-Rab25 cells^32^, while degradation of the ECM is required in invasive migration^15^ to enable cell movement through “ECM barriers^4^”. We observed that MDA-MB-231 cells seeded on 2D collagen I internalised and acidified collagen I while migrating (**Fig. 5a; Supplementary video 2**), suggesting that ECM endocytosis might facilitate cell migration. Consistently, breast cancer cells internalised CDM during migration, and this seemed to occur in the perinuclear region in front of the extending protrusion at the leading edge (**Fig. 5b; Supplementary video 3**). We therefore reasoned that α2β1 integrin and p38 may facilitate invasive migration, by promoting ECM uptake.

To assess this, we treated MDA-MB-231 cells with p38 MAPK inhibitors and found that 10µM and 50µM SB203580 and SB202190 significantly reduced the velocity of MDA-MB-231 cells migrating on CDM, while only 50µM SB203580 and SB202190 impinged on the directionality of cell migration (**Fig. 5c**). Correspondingly, downregulation of MAP3K1, MAPK11, PPP2R1A and α2 integrin significantly reduced the velocity and directionality of MDA-MB-231 cell migration (**Fig. 5d,e**). To confirm the relevance of this process to different cancer types, A2780-Rab25 cells were treated with 50µM SB203580 or siRNAs against MAP3K1, MAPK11, PPP2R1A and α2 integrin and seeded on CDM. Consistently, A2780-Rab25 cell migration was impaired upon p38 inhibition or siRNA downregulation of MAP3K1, MAPK11, PPP2R1A and α2 integrin (**Extended data Fig. 7a-d**). As a WAVE shell complex subunit, PPP2R1A has been shown to modulate migration persistence in the non-transformed mammary cell line MCF10A and MDA-MB-231 cells^43^. To discern if the effect of PPP2R1A on migration required its catalytic activity, cells were treated with okadaic acid. PP2A inhibition significantly reduced the velocity and directionality of cell migration in both MDA-MB-231 and A2780-Rab25 cells, indicating that PP2A phosphatase activity is required for migration (**Fig. 5f; Extended data Fig. 7e**). To study whether regulators of ECM endocytosis similarly affected directional cell migration, we performed a scratch-wound healing assay with a collagen I overlay^48^ in SW1990 cells. We found that α2 integrin and p38 inhibition significantly reduced wound healing closure compared to the control (**Extended data Fig. 7f**). Taken together, these data indicate that MAP3K1, MAPK11, PPP2R1A and α2 integrin, positive regulators of ECM uptake, modulate cell migration, therefore correlating ECM internalisation and cell migration.

**Fig. 7.**
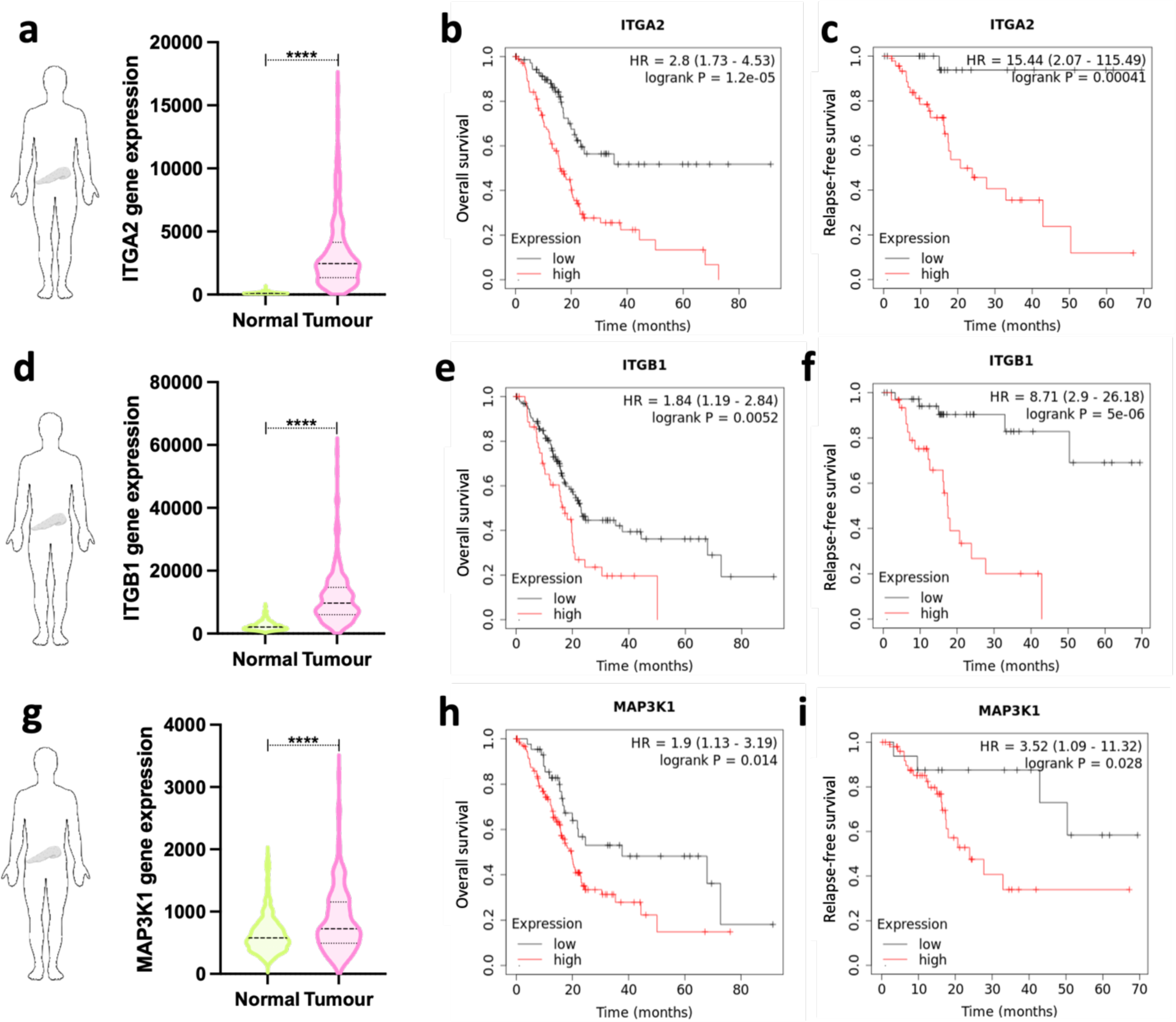
ITGA2 expression was upregulated in mouse mammary cancer and α2, β1 integrin and MAP3K1 expression correlated with poor prognosis of pancreatic ductal adenocarcinoma. **a,d,g,** RNA sequencing data from PDAC tumours (N=177) and normal pancreatic tissue (N=252) for α2 integrin (ITGA2, a); β1 integrin (ITGB1, d) and MAP3K1 (g). ****p <0.0001; Mann-Whitney test. **b,e,h,** Overall survival of patients with PDAC with high (red) or low (black) α2 integrin (1TGA2, b); β1 integrin (ITGB1, e) and MAP3K1 (h) expression. **c,f,i,** Relapse-free survival of patients with PDAC with high (red) or low (black) α2 integrin (ITGA2, c); β1 integrin (ITGB1, f) and MAP3K1 (i) expression.

Since ECM endocytosis is 1) upregulated in breast cancer cells and 2) is required for migration on CDM, we next tested whether regulators of ECM endocytosis were important for 3D invasion (**Fig. 6a**). Indeed, downregulation of MAP3K1 significantly reduced invasion of MDA-MB-231 cells in 3D culture systems (**Fig. 6b,c**). In agreement with the cell migration data, MDA-MB-231 cells were able to internalise ECM in 3D and MAP3K1 knockdown significantly reduced ECM uptake in these settings (**Fig. 6b,d**). We also observed differences in the invasive front morphology, where MAP3K1 knockdown resulted in shorter and thicker multicellular strands protruding from the spheroid core (**Fig. 6b,e**). This observation demonstrates that regulators identified in our 2D screen also play a role in 3D systems, suggesting the screening set up developed in this study has the potential of identifying novel regulators of cancer cell invasion. Similarly to MAP3K1, α2 integrin downregulation or pharmacological inhibition impaired breast cancer cell invasion in 3D systems (**Fig 6f,g**). Altogether, these data demonstrate that MAP3K1 and α2 integrin, regulators of ECM macropinocytosis, are required for MDA-MB-231 cell invasion.

### α2 integrin, β1 integrin and MAP3K1 are poor prognosis factors for pancreatic carcinoma and are linked to chemoresistance in breast cancer

Since ECM endocytosis is upregulated in breast cancer cells (**Extended data Fig.1a,b**), we hypothesised that the ECM uptake regulator α2 integrin might be upregulated during tumour progression *in vivo*. To address this, we stained mammary glands of MMTV-PyMT mice extracted at 44 days (normal mammary gland, preceding tumour formation), 73 days (representing ductal carcinoma in situ (DCIS) stage) and 91 days (invasive adenocarcinoma, IDC). Indeed, α2 integrin expression was slightly higher in DCIS and IDC tissue sections, compared to the healthy tissues (**Extended data Fig. 8a,b**). In breast cancer, chemotherapy resistance is often accompanied by metastatic dissemination, leading to poor clinical prognosis^49^. To examine whether regulators of ECM internalisation may impact on chemoresistance in breast cancer patients, we looked at the correlation between MAP3K1, MAPK11, PPP2R1A and α2 integrin expression and chemotherapy response using transcriptomic data of 3,104 breast cancer patients^50^. Interestingly, these genes were highly expressed in chemoresistant (non-responder) tumours (**Extended data Fig. 8c-f**). Indeed, the area under the ROC curve (AUC) for PPP2R1A and α2 integrin was over 0.6 (**Extended data Fig. 8e,f**), which classified them as a weak biomarker with potential use in prediction of chemotherapy treatment^50^. Given the requirement of α2β1 integrin/p38 MAPK axis in ECM internalisation and invasion in PDAC cells, we looked at the relationship between the expression of α2 integrin, β1 integrin and MAP3K1 and disease outcomes in PDAC patients. RNA sequencing data showed overexpression of α2 integrin, β1 integrin and MAP3K1 in pancreatic adenocarcinomas compared to normal pancreas (**Fig. 7a,d,g**). Correspondingly, high expression of these genes correlated with poor overall survival (**Fig. 7b,e,h**) and relapse-free survival (**Fig. 7c,f,i**). These data indicate that α2 integrin and MAP3K1 may contribute to invasion, metastasis and chemoresistance *in vivo*.

## Discussion

Dysregulations in endocytic trafficking, such as overexpression of trafficking proteins or macropinocytosis, have been associated with cancer^32,51–53^. We showed that Matrigel and CDM internalisation and degradation are upregulated in mouse mammary cancer cells, PyMT#1, compared to normal mouse mammary epithelial cells, NMuMG. These data agree with our previous observations using the MCF10 series of non-transformed, DCIS and invasive cells^5^. Increased lysosomal degradation of CDM, in contrast to Matrigel, in invasive cancer cells may reflect a characteristic acquired during invasion into the surrounding stroma of primary tumours. In fact, starved mouse mammary epithelial cells were reported to internalise and degrade the underlying basement membrane for survival^54^. Following a canonical endocytic route, Matrigel is first delivered into early endosomes and later to lysosomes. This differs from previous results in which fibronectin-occupied α5β1 integrin was directly delivered into lysosomes^32^, suggesting that different ECM components follow distinct trafficking routes. We previously reported that macropinocytosis is the main endocytic route for internalisation of collagen I, Matrigel and CDM^5^. Macropinosome acidification has been shown to occur within 5 to 10 minutes after macropinosome formation^55^, which is in agreement with our live cell imaging observations, where pHrodo-collagen I fluorescence increases ∼ 12 minutes after internalisation.

Integrin signalling has mainly been associated with Src, focal adhesion kinase (FAK) and epidermal growth factor receptor (EGFR) activation, resulting in cell adhesion, proliferation and migration^56–58^. While integrins have been shown to modulate ECM uptake, the signalling regulators downstream of integrin engagement that promote ECM internalisation have remained mainly unexplored. We designed a semi-automated high-content screen and found that MAPK signalling, and in particular p38β, is a key regulator of this process. The role of p38 in tumour development remains controversial. Owing to the high sequence homology between p38α and p38β, few studies have focused on deciphering the role of p38β in tumour progression. Nonetheless, recent evidence proposed that p38β may have distinct and non-redundant functions^59^. The p38 inhibitors used in this study target both p38α (MAPK14) and β (MAPK11)^60^, we thus cannot discard that under certain circumstances MAPK14 may regulate this process. We previously showed that PAK1 modulates macropinocytosis of Matrigel, collagen I, normal fibroblast-and cancer associated fibroblast-CDM^5^. PAK1 is recruited to p38 kinase complex in a phosphorylation-dependent manner^61^. Interestingly, PAK1 has been shown to phosphorylate MAP3K1 on serine 67, which inhibits its binding to JNK kinases^62^, while MAP3K1 has been described as an upstream regulator of MAPK11 and 14^63–66^. 3D collagen I has been shown to lead to p38 activation downstream of α2β1 integrin in mesenchymal cells, such as fibroblasts, in a Cdc42-dependent manner^30^. As PAK1 can be activated by Cdc42, this raises the intriguing hypothesis that PAK1 might promote MAP3K1-dependent p38 activation, leading to ECM uptake, downstream of collagen I binding to α2β1 integrin. Indeed, α2 integrin inhibition or downregulation significantly decreased the uptake of collagen I-rich matrices in MDA-MB–231, A2780-Rab25, YEJ P and SW1990 cells. It is important to note that α2 integrin downregulation had a stronger effect on collagen I than CDM internalisation in A2780-Rab25 cells. This is likely due to the abundance of fibronectin in the CDM, which is internalised in an α5β1-dependent manner in A2780-Rab25 cells ^32^, while fibronectin uptake in MDA-MB-231 cells is very low (data not shown). This may be due to Rab25 not being overexpressed in MDA-MB-231 cells^67^. These results agree with previous studies showing that α2β1 integrin regulates collagen I remodelling through its phagocytosis in fibroblasts^68^. In addition, we have shown that α2β1 integrin trafficks together with internalised collagen I and the downregulation of MAP3K1, MAPK11 or PPP2R1A significantly reduced ECM uptake and the internal pool of α2 integrin. In addition to ECM uptake, we showed that α2 integrin and p38 were also required soluble dextran internalisation. For these experiments, cells were seeded on low collagen I concentrations, which resulted in minimal collagen I uptake (data not shown). It is therefore unlikely that the reduced dextran uptake is due to dextran mostly binding to collagen I and being internalised together with it. Thus, we hypothesise that activation of p38 by α2β1 integrin promotes macropinocytosis of both soluble components, such as dextran, and fibrillar ECM. In fact, p38 inhibition was reported to prevent dextran uptake in dendritic cells^69^, while α5β1 integrins^42^ and α6β1^70^ have been shown to localise to macropinocytic cups and macropinosomes, respectively.

Protein phosphatases have been commonly studied in the context of negative regulation of MAPK signalling pathways; however, our results may indicate that protein kinases (MAP3K1 and MAPK11) may cooperate with PPP2R1A to promote ECM internalisation. Interestingly, the phosphatase activity of PP2A has been shown to dephosphorylate p38 after its activation downstream of collagen I signalling in platelets^29^. Moreover, PP2A is activated downstream of α2β1 integrin in fibroblasts^31^. PP2A is found in integrin adhesion complexes^71^, where it promotes FA maturation and cell migration of the fibrosarcoma cell line HT1080^72^. We propose that MAP3K1 activation downstream of α2β1 integrin promotes MAPK11 activation, with subsequent PP2A activation to regulate the spatiotemporal activation of MAPK11 (Fig. 6h). Interestingly, MAPK11/14 activation by extracellular stimuli inhibits endocytic recycling and promote lysosomal degradation of endocytosed receptors^73^, suggesting that activation of MAPK11/14 may regulate ECM trafficking at different stages. Future studies will focus on the mechanism regulating the trafficking of ECM to the lysosomes.

Integrin internalisation and ECM degradation have been associated with invasive migration and metastatic dissemination. Live cell imaging showed that MDA-MB-231 cells internalise CDM while migrating. Interestingly, pharmacological inhibition or siRNA-mediated downregulation of ECM uptake regulators reduced the velocity and directionality of migrating cells. Of note, 50µM SB202190 and SB203580 affected both the velocity and directionality of migrating MDA-MB-231 cells, in agreement with MAPK11 knockdown. Nonetheless, 10µM SB202190 and SB203580 did not affect the directionality of these cells. This could be in part explained by the fact that the lower p38 inhibitor concentrations had a moderate effect on CDM uptake compared to the higher concentrations. In addition, the IC50 of the compounds for MAPK11 is higher than MAPK14^74^, therefore a higher concentration might be required to fully inhibit MAPK11, which appears to be the major regulator of ECM uptake.

The role of macropinocytosis in modulating invasive migration is controversial, as while α5β1 macropinocytosis decreases invasion in the Ewing sarcoma A-673 cell line^42^, macropinocytosis of α6β1 promotes glioblastoma cell invasion^70^. Interestingly, PPP2R1A is required for embryonic patterning, primitive streak formation, gastrulation and mesoderm formation^75^, which consist of migration processes^76^. This suggests the intriguing possibility that during embryonic development migrating cells remodel the surrounding ECM via macropinocytosis. Indeed, the syncytialisation of placental trophoblasts is accompanied by a strong activation of macropinocytosis^77^. Our results suggest that PP2A activity is required for cell migration of MDA-MB-231 and A2780-Rab25 cells. Indeed, PP2A activity is upregulated in the osteosarcoma cells LM8, MG63 and SaOS cells^78^ and PP2A regulates migration, proliferation and metastasis of osteosarcoma cells^78^. Nevertheless, PP2A was shown to inhibit cervical cancer cell migration by dephosphorylating JNK, p38 and ERK^79^. This suggests that the role of PP2A in cell migration may be cancer-specific. In addition, MAP3K1 downregulation resulted in changes in the multicellular protrusions projecting from 3D spheroids, suggesting either a possible change in the mode of invasion or reduced invasion with limited effects on cell proliferation, resulting in an accumulation of cells in invasive protrusions.

We observed that α2 integrin expression increased in breast and pancreatic tumours and correlated with poor survival. In agreement with our results, patient-derived xenografts derived from breast cancer bone metastasis showed increased α2β1 integrin expression when undergoing epithelial-to-mesenchymal transition with progressive passages through mice^80^. Similarly, α2 integrin expression is induced in ovarian cancer cells that metastasized to the omentum^81^. The role of α2 integrin in pancreatic cancer remains controversial. While α2 integrin expression correlates with poor overall survival and resistance to gemcitabine in high-stiffness matrices in pancreatic cancer^82^, poorly differentiated tumours were reported to express low levels of α2 integrin, while well-differentiated ones showed high levels of α2 integrin^83^. Consistent with our data showing a correlation between MAP3K1 levels and poor prognosis in PDAC patients, MAP3K1 expression correlates with progression and poor prognosis of hormone-receptor-positive, HER2-negative early-stage breast cancer patients^84^. Single-cell transcriptomes from matched primary tumours and metastasis from patient-derived xenograft models of breast cancer showed that metastasis displayed increased stress response signalling during metastatic progression^85^. Interestingly, p38β (MAPK11) activation has been associated with stress signalling^86^, suggesting the intriguing hypothesis that ECM uptake might be induced by the stress response during cancer dissemination. Indeed, MAPK11 promotes metastatic dissemination to the bone, where it promotes osteolytic bone destruction^87^. Similarly, α2 integrin expression promotes bone metastasis of the prostate cancer cell line LNCaP PCa^88^. In addition, integrins and stress responses have been linked to therapy resistance^89^. Indeed, MAPK14 is upregulated in BRAF-inhibition-resistant melanoma in mouse models^89,90^, while MAPK11 abrogation promotes radiosensitivity in A549, MCF7 and HCT-116 cells^91^. Consistently, we showed that MAP3K1, MAPK11, PPP2R1A and α2 integrin expression is higher in chemoresistant breast cancer patients. Dual inhibition of MAPK11 and MAPK14 is currently being tested as a monotherapy or in combination with other agents, such as gemcitabine and carboplatin, for the treatment of glioblastoma, ovarian and metastatic breast cancer. Interestingly, the combination therapy improved progression-free survival in epithelial ovarian cancer^92^. Similarly, α2 integrin inhibitors are being studied in clinical trials for solid tumours, either alone^93^ or in combination with chemotherapy^94^. This raises the intriguing hypothesis that MAPK11/14 activation in therapy-resistant cancers could contribute to metastatic dissemination by promoting ECM-bound α2β1 integrin internalisation. Further work will look into this.

Altogether, we showed that ECM internalisation is upregulated in breast cancer cells and ECM macropinocytosis facilitates invasive migration of breast, ovarian and pancreatic cancer cells, through the activation of p38 signalling. High expression of the regulators of ECM internalisation correlates with poor prognosis of pancreatic cancer patients and is associated with chemoresistance in breast cancer, therefore targeting ECM macropinocytosis could open new avenues for the development of novel strategies to prevent cancer cell invasion and dissemination, which ultimately constitutes the primary cause of cancer death^95^.

## Methods

### Antibodies and reagents

Primary antibodies for Alexa-fluor 488 Anti-human CD29 (ITGB1, Clone TS2/16; 303016), FITC anti-human CD49b (ITGA2; Clone P1E6-C5; 359306), Alexa-fluor 488 anti-mouse CD49b (Clone HMα2; 103510) and mouse anti-human LAMP2 (354303) from BioLegend, primary antibodies for mouse anti-human CD49b (611017) mouse anti-EEA1 (610457) from BD Bioscience. Primary antibody against anti-human GAPDH (Santa Cruz Biotechnology; SC-47724). Secondary antibodies Alexa-fluor 488 Donkey Anti-mouse were from Fisher Scientific (A-21202), IRDye® 800CW and IRDye® 680CW were from LI-COR. Alexa fluor^TM^ 647 Phalloidin, NHS-Fluorescein, NHS-Alexa Fluor 555, pHrodo iFL STP ester red and Hoechst-33342 were from Invitrogen. Collagen I and Matrigel were from Corning. DMEM, RPMI, OptiMEM, Trypsin-EDTA, TrypLE and FBS were from Gibco. E64d (Aloxistatin) was from AdooQ Bioscience. Bafilomcyin A1, FR180204, PD98059, SB202190 and SB203580 from MedChem express. SB203580 from Startech. Okadiac acid from Merk. BTT-3033 from Tocris. siRNA from Dharmacon Horizon discoveries (Singaporean distributor Research Instruments). Vectashield antifade mounting medium from VECTOR laboratories.

### Cell culture

MDA-MB-231 cells, MDA-MB-231 cells overexpressing PPP2R1A-GFP, MDA-MB-231 cells expressing GFP, Telomerase immortalised normal fibroblasts (TIF) and SW1990 cells were cultured in High glucose Dulbecco’s Modified Eagle’s Medium (DMEM) supplemented with 10% fetal bovine serum (FBS). MDA-MB-231 cells overexpressing PPP2R1A were a gift from Professor Alexis Gautreau (École Polytechnique, Paris, France). A2780 overexpressing Rab25, A2780-Rab25, were maintained in Roswell Park Memorial Institute (RPMI) medium supplemented with 10% FBS. NMuMG cells were cultured in DMEM supplemented with 10% FBS and 10µg/ml insulin. PyMT#1 and YEJ P cells were cultured in DMEM 20 ng/ml and 10μg/ml insulin. NMuMG, PyMT#1 and YEJ P cells were a gift from Professor Jim Norman (CRUK Scotland Institute, Glasgow, UK). Cells were grown at 5% CO_2_ and 37◦C and passaged every 3 to 4 days.

### Generation of Cell-derived matrices (CDMs)

CDMs were generated as previously described^96^. CDMs were generated either in a 35mm glass bottom dish, twelve-well plate, 8-well chamber or a 384-well plate. Tissue culture plates were first coated with 0.2% (v/v) gelatin for 1 hour at 37°C. Following that time, plates were washed twice with PBS and crosslinked with 1% (v/v) sterile glutaraldehyde (dissolved in PBS) for 30 minutes at room temperature. Plates were thereafter washed twice with PBS and the remaining glutaraldehyde was quenched with 1M sterile glycine for 20 minutes at room temperature. Subsequently, plates were washed twice with PBS and equilibrated for 30min in complete medium at 37°C. Confluent TIFs were seeded onto the gelatin-coated plates (**Supplementary Table 4**). TIFs were incubated at 37°C in 5% CO_2_ until being fully confluent. The following day or the day after, the media was changed to complete media supplemented with 50µg/ml ascorbic acid, the media was refreshed every other day. TIFs were kept secreting CDM for 9 days in a 10cm dish, twelve-well plate and 35mm glass bottom dish. While only 7 days were required for CDM production in 384-well plates. Following that time, cells were washed once with PBS containing CaCl2 and MgCl2 (PBS++). Cells were incubated with the extraction buffer (20mM NH4OH and 0.5% triton X-100 in PBS++) for 2 to 5 minutes at room temperature until no visible cells remained. For the 384-well plates, cells were extracted twice for 2min. Extracted CDMs were subsequently washed twice with PBS++ and residual DNA was digested with 10µg/ml DNase I in PBS++ at 5% CO_2_, 37°C for 1h; for 384 well plates, DNase incubation was overnight. CDMs were then washed with PBS++ and stored at 4°C in PBS++ supplemented with 1% Penicillin/Streptomycin.

### High-throughput ECM internalisation screen

*<u>Matrigel coating and labelling.</u>* Matrigel preparations were handled using high-grade dispensing pipette tips (E1-ClipTip Equaliser pipette). All the reagents were added in a high throughput fashion by using the multidrop combi at slow or medium speed. The small and large multidrop combi dispenser cassettes, henceforth small or large cassettes, were sterilised with 100ml of 70% ice-cold ethanol. Cassettes were then rinsed with sterile ice-cold water (Gibco). The large cassette dispensed 50µl ice-cold PBS in a 384-well plate. PBS plates were kept at 4°C for 15 minutes. The small cassette was kept at 4°C and was set to dispense 2µl ice-cold Matrigel in the 384-well plates containing PBS. Plates were centrifuged for a few seconds at 500rpm, kept at 4°C for 8 minutes and then polymerised for 2h 30 minutes at room temperature. Bravo liquid handling system (Agilent Technologies, hereafter Bravo) was used to perform Matrigel washes. After polymerisation time was completed, 40µl PBS was pipetted up and deposited into the waste reservoir. The multidrop combi large cassette was set to dispense 30µl/well of 20µg/ml pHrodo in a 384-well plate. Plates were kept in the dark on gentle rocking for 1h for efficient Matrigel labelling. Following that time, Bravo was used for washing pHrodo off. All washes were performed with PBS containing 1% antibiotic-antimycotic (anti/anti) to avoid contamination. Plates were kept in 50µl 1% anti/anti in PBS at 37°C overnight. *<u>Cell detachment.</u>* VIAFLO-384-well head (Integra) was used for cell transfer. Cell media in transfected plates was automatically pipetted and released in a waste reservoir. Transfected plates were washed with 80µl PBS once and 20µl of TrypLE was used to ensure the detachment of cells. Cells were incubated for 5 minutes at 37°C. Cells were then vortexed every 1 minute thrice. TrypLE^TM^Express Enzyme was neutralised with 80µl 10%FBS DMEM. Cells were pipetted up and down before transferring to ECM-coated 384 well plates. Transfected and transferred cells were incubated for 6 hours. After 2 hours, cell media was changed to 200nM Bafilomycin A1 in positive control wells containing NT5. After 5h 45 minutes, Hoechst-33342 was spiked into a final concentration of 0.5 to 1µg/ml. Cells were imaged with 40X water objective Opera Phenix. The screen was performed in duplicate; the same imaging and analysis threshold was applied for both replicates. Data was normalised between the average of the non-targeting (NT5) control and NT5 in the presence of bafilomycin A1, as a positive control to assess the magnitude of modulation between technical replicates, similar to^97^. To identify the hits, we calculated the first derivative of the curve of normalised values within the population tested, excluding controls. We identified 25 and 37 positive regulators, and 66 and 55 negative regulators for the first and second replicate, respectively. To rule out that changes in ECM uptake were a consequence of changes in apoptosis, survival and/or proliferation, we assessed the nuclei count and considered a nuclei count lower than 200 as an indication of cell toxicity of the knockdown as in^23^.

### ECM internalisation

Collagen I were dissolved in ice-cold PBS to a final concentration of 1mg/ml. 100μl of the solution was used to coat a 35mm glass-bottom dish with the help of a pipette tip. The ECM-coated dishes were incubated at 37°C and 5% CO_2_ for 1h for polymerisation. Collagen I dishes were labelled with either 300µl of 10µg/ml NHS-fluorescein, 5µg/ml Alexa fluor™555 NHS ester or 20µg/ml pHrodo iFL red. The dishes were incubated for 1 hour at room temperature on gentle rocking. CDMs were labelled with 20µg/ml pHrodo iFL red for 1h at 37°C. The labelled ECM dishes were washed twice with PBS prior to cell seeding. To avoid evaporation, PBS was added to each dish to keep in the incubator. Alternatively, CDMs were labelled with 0.13mg/ml NHS-SS-biotin in PBS++ for 30min with gentle rocking at 4°C, labelled CDMs were washed twice and cells were seeded on top. After the incubation time, cells were washed once with ice-cold PBS++ and treated with a cell-impermeable reducing agent (15mg/ml sodium 2-mercaptoethane sulfonate supplemented in 3mM NaOH for 1h 30min at 4°C). Reduced cysteines were alkylated with 17mg/ml iodoacetamide for 10min at 4°C. Dishes were kept on ice, washed once with PBS++ and fixed with 4% formaldehyde. To detect uptake in these cells, cells were permeabilised and stained with Streptavidin Alexa fluor 488 (1:1000) at room temperature for 1h. For pHrodo-labelled ECM, cells were stained with Hoechst-33342 and imaged live. For p38 MAPK inhibitors, cells were serum starved for 16 to 18h. Cells were detached using TrypLE and neutralised in serum-free media. Preliminary screening of MAPK inhibitors was done in serum-free conditions to assess ECM-specific contribution and avoid MAPK activation by growth factors found in serum^30^; other assays were performed in 5% FBS. For uptake, cells were incubated for 6h, except for YEJ P cells, which were kept for 16h. For BTT-3033 experiments, cells were allowed to adhere for 2h before adding 5µM or 10µM BTT-3033 depending on the cell line. MDA-MB-231 were cultured for an additional period of 6h in the presence of BTT-3033 before fixation and immunofluorescence staining. SW1990 cells were cultured for 4 additional hours before nuclei labelling and live imaging. For knockdown experiments, cells were incubated for a total of 6h on the different types of ECM. For time-lapse uptake, MDA-MB-231 cells were seeded on collagen I matrices labelled with NHS-fluorescein and pHrodo red. After 30 minutes of cell seeding, cells were imaged every minute with a 63X oil objective Zeiss LSM980 Airyscan 2 microscope for a total period of 5 to 6h.

### Rhodamine-dextran internalisation

MDA-MB-231 cells were seeded at a density of 10^5^ cells/well for 6h on 0.1mg/ml collagen I (50µl/well)-coated 8-well chamber. Then, cells were pre-treated with DMSO or 50µM SB202190 for 30 minutes. Cells were later incubated for 1 hour with 0.25mg/ml Rhodamine-dextran in the presence of SB202190 or DMSO. Alternatively, MDA-MB-231 cells were seeded at a density of 3×10^5^ cells/dish on 35mm glass-bottom dishes coated with 0.1mg/ml collagen I for 5h in the presence of BTT-3033 or DMSO. Cells were later incubated for 1 hour with 0.25mg/ml Rhodamine-dextran in the presence of the inhibitors or the vehicle control. Cells were then fixed with 4% paraformaldehyde and stained for human β1 integrin (1:400 dilution). Vectashield mounting medium with DAPI allowed the visualisation of the nuclei. Cell imaging was carried out with a 60x objective Nikon A1 confocal microscope. Image J^98^ was used to analyse Dextran uptake index^99^.

### Immunofluorescence

Cells were fixed with 4% (w/v) formaldehyde in PBS for 15 minutes. Next, cells were permeabilised with 0.25% (v/v) Triton X-100 in PBS for 5min and washed twice with PBS. For dextran internalisation assays, cells were not permeabilised. For ECM internalisation assays, cells were stained for the actin cytoskeleton with Phalloidin conjugated with either Alexa Fluor 555. Phalloin was diluted 1:500 in PBS; cells were incubated for 10 minutes at room temperature. For antibody staining, cells were blocked in 1% (w/v) bovine serum albumin (BSA) for 1 hour at room temperature. Cells were then incubated with the respective primary antibodies for 1 hour at room temperature. Anti-human ITGB1 antibody conjugated to Alexa fluor 488 (1:400 in PBS) was used for the colocalization experiment. For α2 integrin staining, cells were incubated in FITC anti-human CD49b antibody (1:200 dilution) in PBS. For non-fluorescently conjugated antibodies, EEA1 and LAMP2, cells were first incubated with the primary antibody (1:100 in PBS) for 1h at room temperature. Cells were then washed thrice with PBS and incubated with the secondary antibody Alexa-fluor 488 Donkey Anti-mouse (1:1000) for 45 minutes at room temperature. Cells were washed three times with PBS and once with ionised water. Vectashield antifade mounting media with DAPI was used for nuclear staining and sample preservation. For colocalization experiments, the ‘colocalization colormap’ ImageJ plug-in was used^100^.

### siRNA transfection

For 384-well plates, 2.5µl of 500nM siGENOME siRNA and 2.5µl Opti-MEM per well were added into CellCarrier Ultra 384-well plates (Perkin Elmer). 4.95µl Opti-MEM was incubated with 0.05µl Dharmafect IV for 5 minutes. 5µl of the Dharmafect IV solution was added into each well. Plates were incubated for 20 minutes on gentle rocker at RT. 3×103 cells were seeded in 40µl DMEM containing 10% FBS. The final concentration of the siRNA was 25nM. Cells were kept at 5% CO_2_ and 37°C for 72h. β1 integrin knockdown efficiency was analysed in Columbus software. For 6-well plates, 10µl 5µM siRNA (**Supplementary Table 5**) were mixed with 190µl Opti-MEM into each well of a 6-well plate. 198µl Opti-MEM and 2µl Dharmafect I were mixed and incubated for 5 minutes at RT. 200µl of the Opti-MEM Dharmafect I mix was added on top of the siRNA and incubated for 20 minutes on a rocker. 4×105 cells in 1.6ml were added into each well. Cells were incubated at 37°C and 5% CO_2_ for 72h. Following this time, cells were used for uptake experiments or analysis of knockdown efficiency by western blot or RT-qPCR. For PPP2R1A knockdown, cells were seeded on a glass bottom dish and knockdown efficiency was analysed in ImageJ. Alternatively, for ITGA2 migration and invasion assays in MDA-MB-231 cells, cells were seeded in 6-well plates. The next day, 5µl of Lipofectamine-2000 were mixed in 250µl of Opti-MEM (Solution A) and 5µl of 20µM siRNA were mixed with 250µl Opti-MEM (Solution B). Both solutions were mixed and incubated for 20 minutes at room temperature. Plates containing adhered cells were washed once with PBS and 500µl of Solution A+B was added on top of each well, 500μl of Optimem was added and cells were incubated for a period of 4 to 6h. After this time, the media was aspirated and 2ml of fresh 10% DMEM was added; alternatively, for 3D spheroid generation, knockdown cells were cultured into methylcellulose drops for 48h.

### Western blotting

Confluent 6-well plates were harvested with 100µl of lysis buffer (50mM Tris-HCl pH 7 and 1% SDS). Cell lysates were collected and transferred into QiaShredder columns (Qiagen), which were spun at 4°C for 10 minutes at 13000rpm. Extracted proteins were mixed in a 4:1 ratio with the NuPAGE buffer with a final concentration of 1mM DTT. 15µl to 25µl of extracted proteins in NuPAGE buffer and 0.5µl protein ladder (BioLabs) were loaded into a Bio-Rad 4-15% Mini-PROTEAN precast polyacrylamide gel. The gels were run at 100V constant voltage for 1h 15 minutes. Running buffer: 3g Tris base, 14.4g glycine and 1g SDS in 1l ionised water. Afterwards, proteins were transferred to a FL-PVDF membrane using the Towbin transfer buffer (25mM Tris, 192mM glycine, 20% methanol (v/v) pH 8.3). Membrane transfer was performed at room temperature, constant voltage 100V for 1h 15 minutes. Membranes were blocked in 5% milk for 1 hour at room temperature. Membranes were washed twice TBST (50mM Tris HCl, 150mM NaCl and 0.5% (w/v) Tween 20), and incubated 1h 30min with the primary antibody anti-human CD49b and mouse anti-human GAPDH in TBST. Membranes were washed thrice in TBST (10 min/wash). Secondary antibodies were then applied for 1h at room temperature. Anti-mouse IgG secondary LiCOR IR Dye 800 antibody was diluted 1:30000 in 0.01% (w/v) SDS TBST. Next, three TBST washes were performed and the last wash with deionized water. A LiCOR Odyssey Sa system was used for imaging the membranes. The intensity of the bands was quantified with Image Studio Lite software. Bands were normalised to GAPDH intensity.

### Cell migration

For p38 inhibitor experiments, cells were serum starved for 16 to 18h. Cells were detached using TrypLE and neutralised with serum-free media. 5×10^4^ MDA-MB-231 cells per well were seeded into a CDM-coated 12-well plate. Migration experiments were carried out with 5% FBS DMEM. DMSO, p38 MAPK inhibitors (10µM and 50µM SB202190 and SB203580) and PP2A inhibitor (50nM okadaic acid) were added at the moment of cell seeding. Time-lapse imaging was done after a 6h-incubation. For knockdown experiments, cells were plated in complete media; cells were allowed to adhere for 6h before imaging. Plates were imaged in a Nikon Inverted Ti eclipse with Oko-lab environmental control chamber with a 10X/NA 0.45 objective. Cells were incubated at 37°C and 5% CO_2_; images were acquired every 10 minutes for at least 7h and more than 40 cells per well were quantified per biological replicate. Individual cell migration was manually tracked using MTrack2, a plugin found in ImageJ. The chemotaxis tool plugin in Image J (https://ibidi.com/chemotaxis-analysis/171-chemotaxis-and-migration-tool.html) was used to calculate the velocity and directionality of migrating cells.

### Wound healing

For wound healing experiments, SW1990 cells were detached using Trypsin-EDTA and neutralised using 10% FBS DMEM. Cells were seeded into a 12-well plate and incubated overnight to achieve confluency. Monolayers were manually scratched into a cross shape with the help of a 200μl pipette tip. Cells were covered by 0.5mg/ml collagen I in 10%FBS DMEM (300μl/well) and incubated for 30 minutes at 37°C to polymerise. After this time, 1ml media was added on top of the scratches in the presence of the vehicle (DMSO), 10μM BTT-3033 or 50μM SB203580. Each branch of the scratches was imaged at 0h, 3h and 6h after adding the inhibitors using an Olympus Inverted Fluorescence Microscope (4X objective). The area of the scratch was calculated by tracing the perimeter of the non-invaded area using Image J. The invasion ratio was normalised to the area at time 0h.

### mRNA expression (RT-qPCR)

mRNA was extracted from snap-frozen samples, and stored at −80°C, according to the manufacturer’s protocol (RNeasy® Mini – Qiagen). For one-way qPCR (analysis of MAP3K1, Hs_MAP3K1_1_SG QuantiTect Primer Assay, Qiagen); Luna® Universal One-Step RT-qPCR Kit was used. Samples were prepared by adding less than 1µg RNA, 5µl Luna Universal One-Step Reaction Mix, 0.5µl Luna WarmStart® RT Enzyme Mix, 0.8 mix forward and reverse primer and toped up with RNase-free water for a total of 10µl. For two-way qPCR (analysis of MAPK11; Hs_MAPK11_1_SG QuantiTect Primer Assay, Qiagen); cDNA was first synthesised with High Capacity cDNA Reverse Transcription Kit (Fisher). Afterwards, loading master mix containing 5µl QuantiNova SYBR® Green PCR Kit (Qiagen) master mix, 1µl forward and reverse primer and 1µl RNase free water was prepped and mixed with 3ul cDNA solution (5ng/µl). Finally, for both methodologies used, 10µl of the sample was loaded into a 384-well plate. Quantstudio 12K flex real-time PCR system was used in the SYBR® mode to analyse the samples. Expression levels were calculated using the 2”DDCt method ^101^. GAPDH was used as a control (Hs_Gapdh_3_SG QuantiTect Primer Assay, Qiagen). Each sample was tested in three technical replicates.

### DNA transfection

MDA-MB-231 stably expressing GFP were generated as described in^5^. Briefly, 8×10^5^cells/well were seeded into a 6-well plate in 2ml of 10% FBS DMEM without antibiotics. Confluent cells were transfected with 2.5µg of pSBtet-GB GFP Luciferase plasmid and 0.25µg of the sleeping beauty transposon plasmid, pCMV(CAT)T7-SB100. 250µl of OptiMEM, 5µl p3000 and 3.75µl Lipofectamine 3000, together with both plasmids were added on top of the 2ml. Media was changed after 6h. Following 48h, cells were selected with 2µg/ml blasticidin. After selection, cells were FACS sorted.

### 3D ECM internalisation and 3D spheroids

3D spheroids were generated by the hanging drop method, previously described in ^102^. 2000 cells per 20μl drop containing 4.8mg/ml methylcellulose (Sigma-Aldrich) and 20μg/ml soluble collagen I (BioEngineering) were pipetted on the lid of tissue culture dishes. Lids were turned and put on top of the bottom reservoir of the dish, which was filled with PBS to prevent evaporation. After 48 hr, spheroids were embedded in 40μl of 3mg/ml rat tail collagen I (Corning) and 3mg/ml Matrigel (Corning). For 3D uptake assays, ⅕ (v/v) of the matrix solution was labelled with a final concentration 20µg/ml pHrodo containing 0.1M sodium bicarbonate. Cells were imaged live every 24h until day 3 post-embedding for α2 integrin knockdown and BTT-3033 treatment. For MAP3K1 knockdown, spheroids were imaged until day 4. Nikon A1 confocal (10X objective) was used to image whole spheroids. For invasion analysis, spheroids were threshold and the area of the selected threshold was quantified in Image J. The invasion rate was defined as the ratio between the invasion area and the total area. For 3D uptake, the pHrodo-ECM intensity in the spheroid was quantified.

### Overall survival, relapse-free survival and ROC analysis

The survival analysis was performed using Kaplan-Meier plotter (https://kmplot.com/analysis/), which can assess the effect of genes of interest on survival in 21 cancer types, including pancreatic cancer. Sources for the databases include Gene Expression Omnibus (GEO), European Genome-Phenome Archive (EGA), and The Cancer Genome Atlas (TCGA)^103^. RNA sequencing data from pancreatic ductal adenocarcinoma and normal pancreas was performed using TNMplot (tnmplot.com), which enables a direct comparison of tumour and normal samples and runs a Mann-Whitney U test ^104^. The ROC plotter was used to analyse the link between gene expression and response to chemotherapy (including, taxane, anthracycline, ixabepilone, CMF, FAC and FEC) using transcriptome-level data of breast cancer patients (https://www.rocplot.com/)^50^. Breast cancer datasets were identified in GEO (https://www.ncbi.nlm.nih.gov/gds), using the platform IDs “GPL96”, “GPL570”, “GPL571” and the keywords “breast”, “cancer” and “therapy”. For genes with multiple probes, Jetset was used to select the most reliable probe set (https://services.healthtech.dtu.dk/services/jetset/).

### Immunofluorescence of FFPE tissue samples

Samples were provided by SEARCHBreast (https://searchbreast.org/). Tissue slides were deparaffinized and rehydrated. Antigen retrieval with sodium citrate buffer at pH 6, blocked for endogenous peroxidase with 3% hydrogen peroxide, permeabilised with 1% FBS, 0.5% Triton X-100 in PBS and blocked for non-specific binding with 5% FBS 0.5% Triton X-100 in PBS. Staining for α2 integrin (Alexa-fluor 488 anti-mouse CD49b, 1:200 in 5% FBS, 3% Bovine serum albumin, 0.5% Triton X-100 in PBS) was performed overnight. Slides were counterstained with Hoechst-33342 and then mounted (ProLong™ Gold Antifade Mountant) before fluorescence imaging with Nikon Inverted Ti eclipse with a 10X/NA 0.45 objective. The mean intensity of the mammary glands or tumour cells from the tissue sections was quantified using ImageJ.

### Statistical analysis

All the gathered data was normalised to the control population. Data representation and statistical analysis were performed in GraphPad Prism (Version 9.4.1) software. Scatter plot data are represented by SuperPlots, which enables the incorporation of cell-level data and experimental repeatability in a single diagram^105^. Cell-level data is represented by dots. For images acquired at Nikon A1 confocal, cell data is colour-coded in blue, red and green to display individual biological replicates. SuperPlots in addition include the average (mean) data in each biological replicate for Nikon A1 confocal data. Average values or well data is represented by squares. For images acquired in Opera Phenix microscope, blue, red and green squares show mean biological replicates. Multiple squares with the same colour represent technical replicates for each well. Data acquired was analysed at the cell level. To compare two datasets, an unpaired t-test was used; to compare more than two datasets, nonparametric one-way ANOVA was performed. High throughput screening data was normalised between the NT5 and NT5 in the presence of Bafilomycin A1 (Normalised index = (Matrigel uptake index - mean NT5)/(Mean NT5-Mean Bafilomycin A1)).

## Acknowledgements

Imaging work was performed at the Wolfson Light Microscopy Facility, University of Sheffield, using the Nikon A1 confocal, Nikon widefield and Airyscan microscope. High-throughput imaging was performed in the RNAi screening facility in IMCB, Singapore. FACS sorting was performed by SIgN (A*STAR, Singapore). qPCR analysis was performed in collaboration with the Tsakiridis lab at the University of Sheffield. We would like to acknowledge Marga Albu for developing the Image J macro used for ECM uptake quantification. We thank Dr Rebecca Bennion for the critical reading and proofreading of the manuscript. E.R. is funded by CRUK (C52879/A29144) and this work has also been supported by the Academy of Medical Sciences, Wellcome Trust, Government Department of Business, Energy and Industrial Strategy and British Heart Fundation, Springboard Award (SBF003\1045). M.L.M. is funded by Sheffield/ARAP PhD program. The Wolfson Light Microscopy Facility, University of Sheffield, is funded by the Wellcome Trust (grant WT093134AIA).

## Author contributions

MLM and ER conceived, designed the experiments and interpreted the results. ER, FB and XLG supervised the study. MLM and XLG conceived and developed experimental tools for high-content screening. MLM performed in vitro uptake and migration experiments, developed uptake in 3D spheroid assays and performed tissue staining. KN designed and performed colocalization, collagen I and CDM uptake, migration and spheroid invasion experiments (Fig. 2 a,b; Fig. 5e; Fig. 6f,g; Extended Data Fig. 1a,b,c,d and Extended Data Fig. 3b). ZB performed collagen I, dextran uptake and western blot experiments (Fig. 4f, Extended Data Fig. 3g, Extended Data Fig. 5i-k and Extended data Fig. 6c,d). RB ran qPCR samples, performed and analysed wound healing migration assay (Extended Data Fig. 5e,f and Extended Data Fig. 7f). SY ran qPCR samples and performed migration experiments (Extended Data Fig. 5g,h and Extended Data Fig. 7e). RB and SY wrote the corresponding method section. XLG provided conceptual input and expertise in high throughput screening and liquid handling. JT performed the time-lapse imaging for collagen I uptake (Fig. 5a and Extended Data Fig. 1e). ER and FB acquired funding for this study. MLM wrote the manuscript. ER and XLG provided input on data presentation. MLM and ER edited and reviewed the manuscript. All authors read and approved it.

## Competing interests

The authors declare no competing interests.

---

**Extended data Fig. 1.**
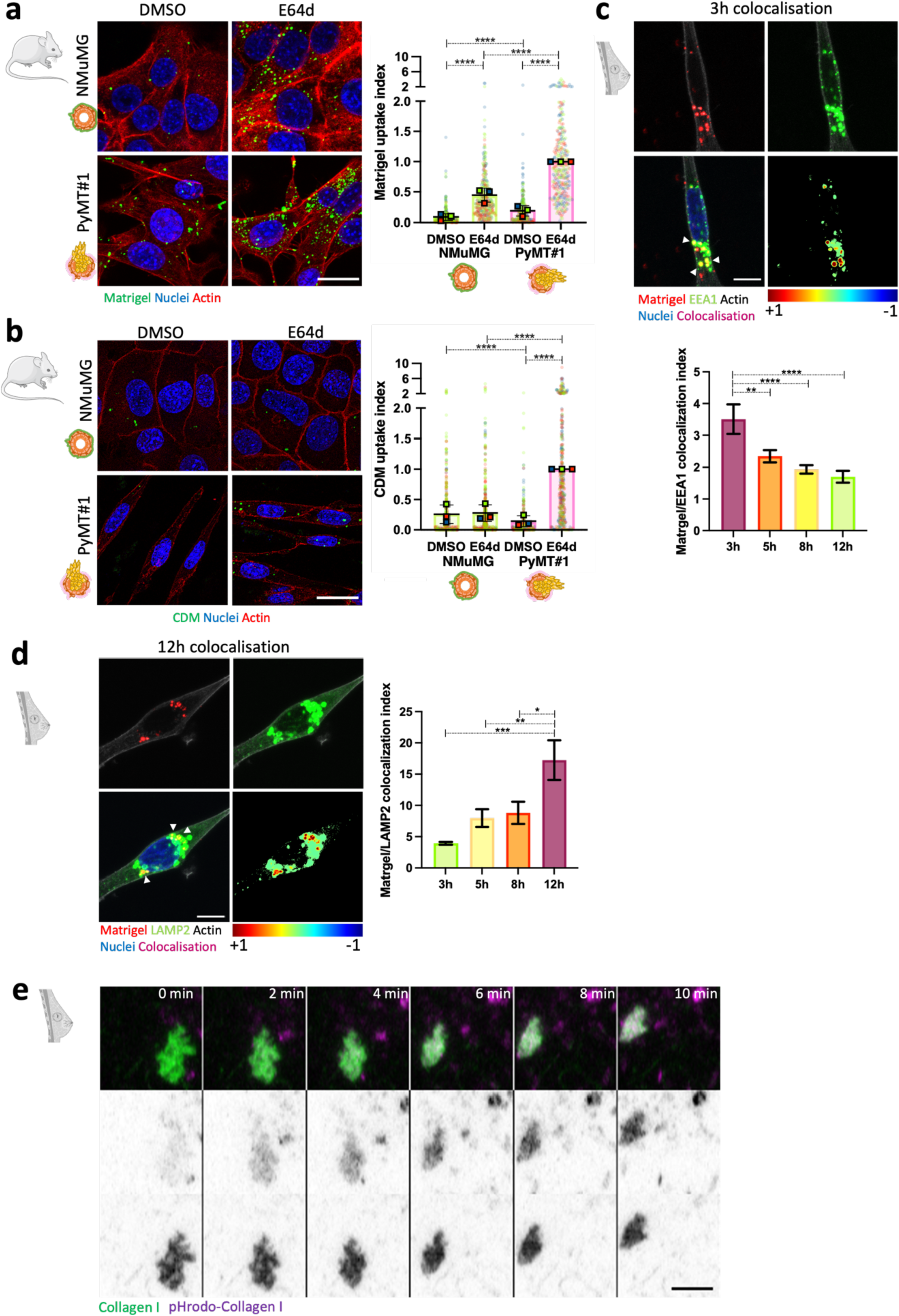
The ECM was delivered to and degraded in the lysosomes. **a,** NMuMG and PyMT#1 cells were seeded on NHS Fluorescein-labelled 1mg/ml matrigel for 12h in the presence or absence of 20µM E64d, fixed, stained for actin and nuclei and imaged with a Nikon Al confocal microscope. Scale bar, 20µm. Matrigel uptake index was calculated with Image J. Data are presented as the normalised mean ± SD; N=3 independent experiments. ****p<0.0001; Kruskal-Wallis test, b, NMuMG and PyMT#1 cells were seeded on biotinylated CDM for 12h in the presence or absence of 20µM E64d, fixed, stained with streptavidin Alexa Fluor 488, Phalloidin Alexa Fluor 555 and DAPI, imaged and quantified as in a. Scale bar, 20µm. Data are presented as the normalised mean + SD; N=3 independent experiments. ****p<0.0001; Kruskal-Wallis test. **c,d,** MDA-MB-231 cells were seeded on NHS Alexa Fluor 555-labelled 1mg/ml matrigel for 3h, 5h, 8h and 12h, fixed, stained for EEA1 (c) or LAMP2 (d), actin and nuclei and imaged as in a. Scale bar, 10μm. Colocalisation was quantified with Image J. Data are presented as the mean ± SEM; N=3 independent experiments. **p=0.0057,****p<0.0001 (c); *p=0.0207, **p=0.0069,***p=0.0003 (d); One-way ANOVA/Tuke/s multiple comparisons test, **e,** MDA MB-231 cells were seeded on NHS-fluorescein (green) and pHrodo-labelled (magenta) 1mg/ml collagen I and imaged live for 5h. Representative time frames from supplementary video 1 are shown. Scale bar, 5µm.

**Extended data Fig. 2.**
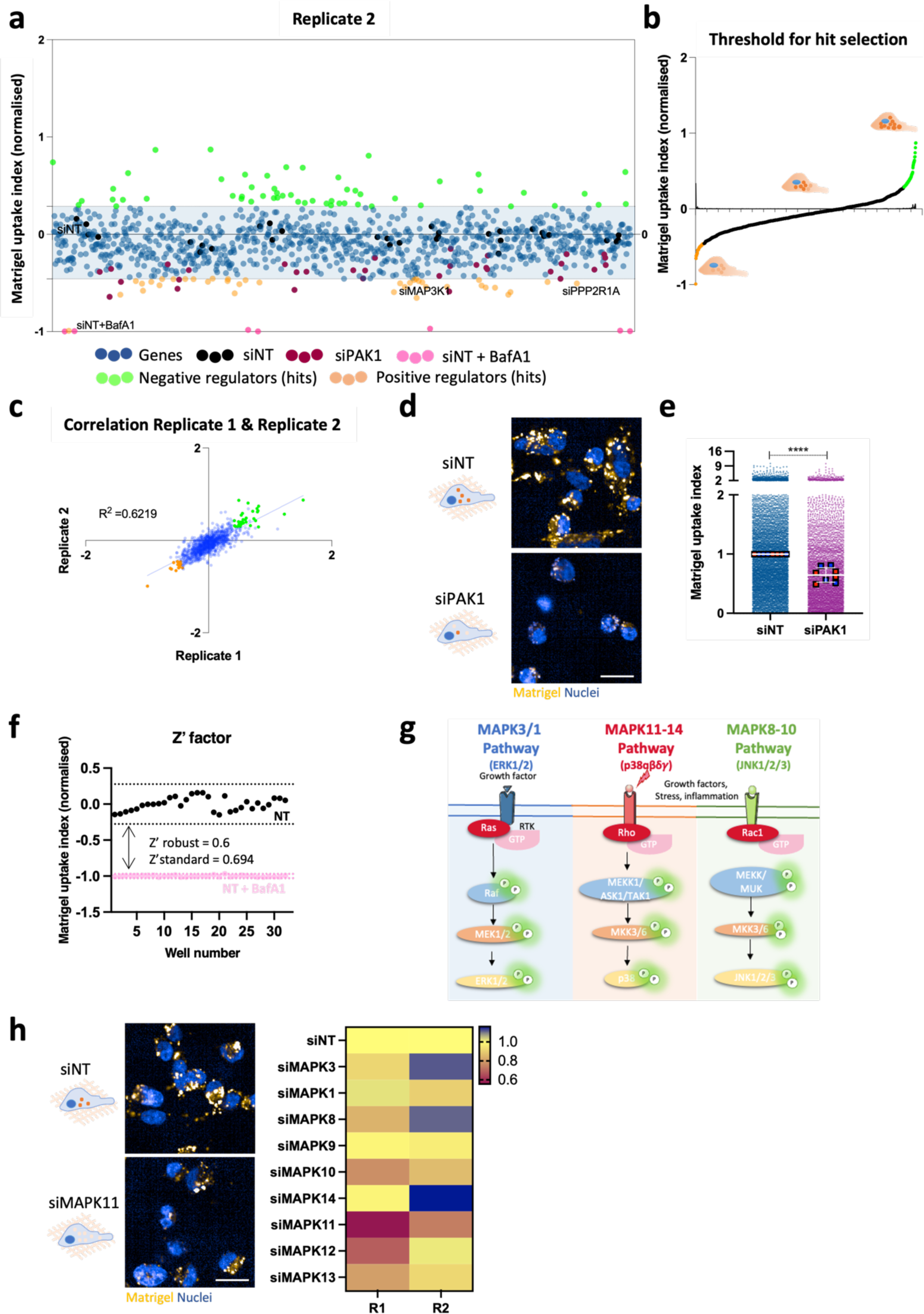
A kinome and phosphatome screen identified regulators of matrigel internalisation. **a,** Normalised cloud plot analysis from replicate 2. **b,** First derivative of the curve for hit threshold for replicate 2. **c,** Correlation between replicate 1 and 2. d. Representative images of matrigel uptake in cells transfected with a non-targeting siRNA control (siNT) and an siRNA targeting PAK1 (siPAK1). Scale bar, 20µm. **e,** Data are presented as the normalised mean ± SD; N=6 replicates from 2 independent experiments. ****p<0.0001; Kruskal-Wallis test, **f,** Z’ robust and standard for screening validation, **g,** Diagram of MAPK activation pathways, **h,** Heatmap of major MAPKs in the kinome and phosphatome screen. N=2 biological replicates. Representative images for MAPK11. Scale bar, 20µm.

**Extended data Fig. 3.**
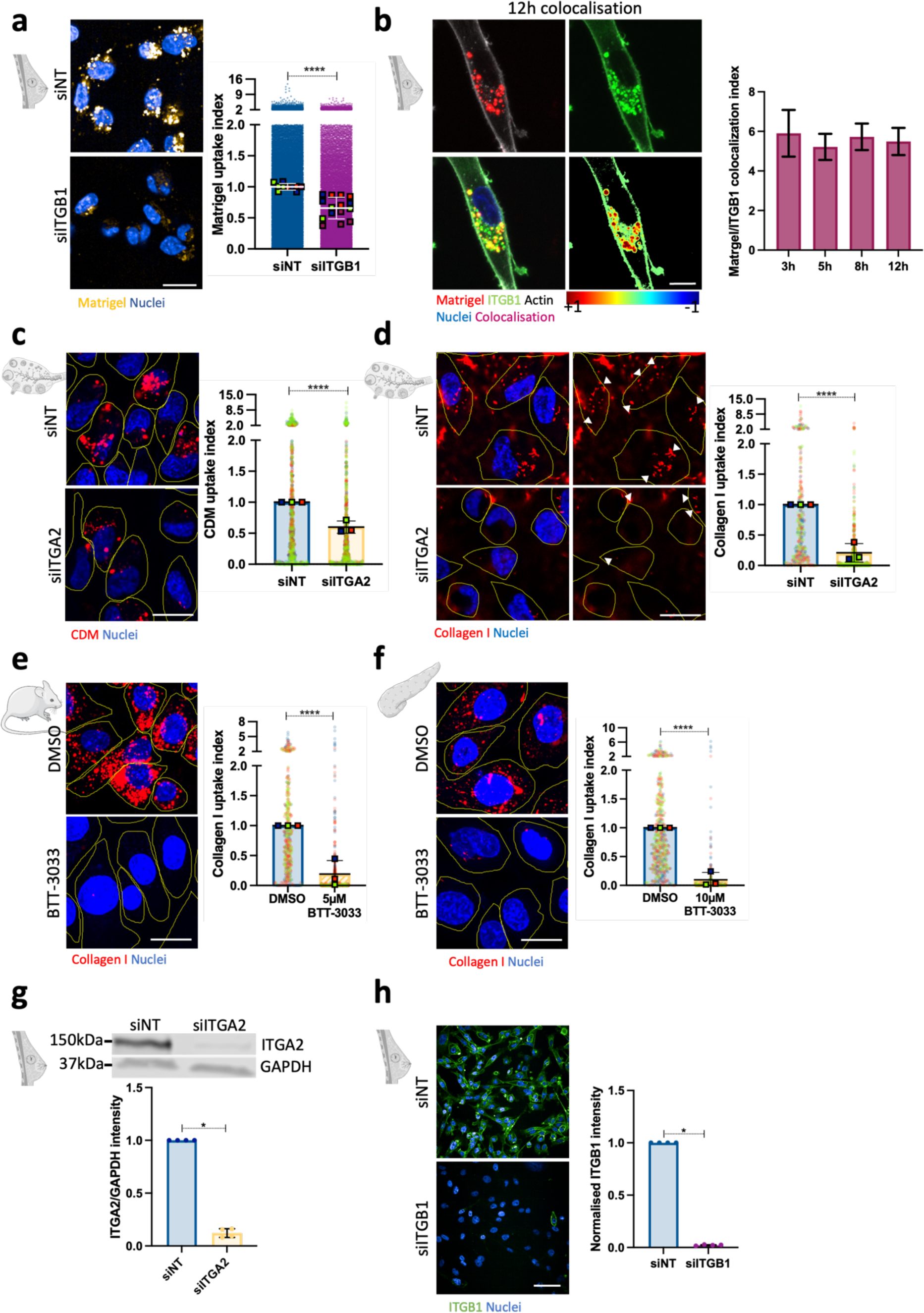
α2 and β1 integrin mediated ECM internalisation in cancer cells. **a,** MDA-MB-231 cells transfected with an siRNA targeting β1 integrin (silTGB1) or a non-targeting siRNA control (siNT), plated on pHrodo-labelled 0.5mg/ml matrigel for 6h, stained with lμg/ml Hoechst and imaged live. Data are presented as the normalised mean ± SD; N=3 independent experiments. ****p<0.0001; Mann-Whitney test, **b,** MDA-MB-231 cells were seeded on NHS Alexa Fluor 555-labelled 1mg/ml matrigel for 3h, 5h, 8h and 12h, fixed and stained for β1 integrin (ITGB1), actin and nuclei. Scale bar, 10μm. Data are presented as the mean ± SEM; N=3 independent experiments. **c,d,** A2780-Rab25 cells were transfected with an siRNA targeting α2 integrin (silTGA2) or a non-targeting siRNA control (siNT), seeded on pHrodo-labelled CDM (c) or collagen I (d) for 6h, stained with 1μg/ml Hoechst and imaged live. Data are presented as the normalised mean ± SD; N=3 independent experiments. ****p<0.0001; Mann-Whitney test, **e,** YEJ P cells were allowed to adhere to pHrodo-labelled 1mg/ml collagen I for 2h, treated with 5μM BTT-3033 or DMSO for 14h, stained with 1μg/ml Hoechst and imaged live. Data are presented as the normalised mean ± SD; N=3 independent experiments. ****p<0.0001; Mann-Whitney test, **f,** SW1990 cells were seeded on pHrodo-labelled 1μg/ml collagen I for 2h, treated with 10μM BTT-3033 or DMSO for 4h, stained with 1μg/ml Hoechst and imaged live. Data are presented as the normalised mean ± SD; N=3 independent experiments. ****p<0.0001; Mann-Whitney test, **g,** Cells were transfected with an siRNA targeting α2 integrin (silTGA2) or a non-targeting siRNA control (siNT) for 72h, lysed and α2 integrin and GAPDH protein levels were measured by western blotting. Data are presented as the normalised mean ± SD; N=4 independent replicates. *p=0.0286; Mann-Whitney test, **h,** Cells were transfected with an siRNA targeting pi integrin (silTGB1) or a non-targeting siRNA control (siNT) for 72h, fixed and stained for pi integrin (ITGB1) and nuclei. Images were acquired with a 40X Opera Phenix microscope. Scale bar, 60um. Data are presented as the normalised well mean ±SD; N=4 independent experiments. *p=0.0286; Mann-Whitney test.

**Extended data Fig. 4.**
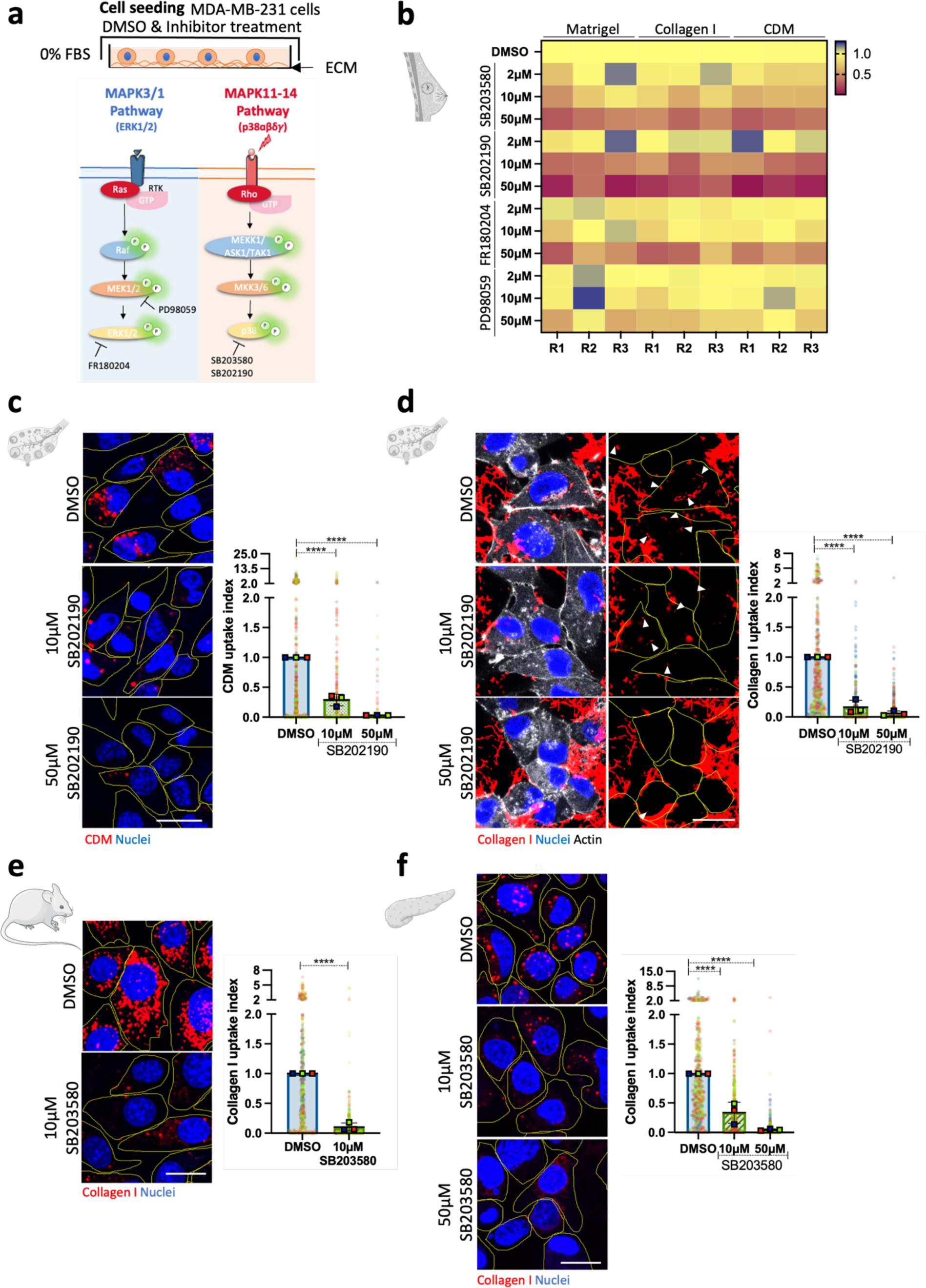
p38 MARK inhibition reduced ECM internalisation. **a,** Schematic representation of MAPK inhibitors, **b,** MDA-MB-231 cells were serum starved for 16 to 18h. 10^4^ cells were seeded on pHrodo-labelled ECM for 6hr in the presence of DMSO or MAPK inhibitors in 0% FBS, stained with 1μg/ml Hoechst and imaged live. Data analysis was performed with Columbus software, **c,** A2780-Rab25 cells were serum starved for 16 to 18h. 3×l0^5^ cells were seeded on pHrodo-labelled CDM for 6h in the presence of DMSO, 10μM or 50μM SB202190 in 5% FBS, stained with Hoechst and imaged live. Scale bar, 20μm. CDM uptake was quantified with ImageJ. Values represented are normalised mean + SD from N=3 independent experiments; ****p<0.0001; Kruskal-Wallis test, **d,** A2780-Rab25 cells were serum starved for 16 to 18h. 3×l0^5^ cells were cultured on 1mg/ml collagen I, labelled with NHS-Alexa fluor 555, for 6h in the presence of DMSO, 10μM or 50μM SB202190 in 5% FBS. Cells were fixed and stained for actin and nuclei. Scale bar, 20μm. Collagen I uptake was quantified with ImageJ. Values represented are normalised mean + SD from N=3 independent experiments; ****p<0.0001; Kruskal-Wallis test, **e,** YEJ P cells were serum starved for 18h. 3×l0^5^ cells were seeded on pHrodo-labelled 1mg/ml collagen I for 16h in the presence of DMSO or lOpM SB203580 in 5% FBS, stained with Hoechst and imaged live. Scale bar, 20μm. Collagen I uptake was quantified with ImageJ. Values represented are normalised mean + SD from N=3 independent experiments; ****p<0.0001; Mann-Whitney test, **f,** SW1990 cells were serum starved for 18h. 3×10^5^ cells were cultured on pHrodo-labelled 1mg/ml collagen **I** for 6h in the presence of DMSO, lOpM or 50pM SB203580 in 5% FBS, stained with Hoechst and imaged live. Scale bar, 20μm. Collagen I uptake was quantified with ImageJ. Values represented are normalised mean + SD from N=3 independent experiments; ****p<0.0001; Kruskal-Wallis test.

**Extended data Fig. 5.**
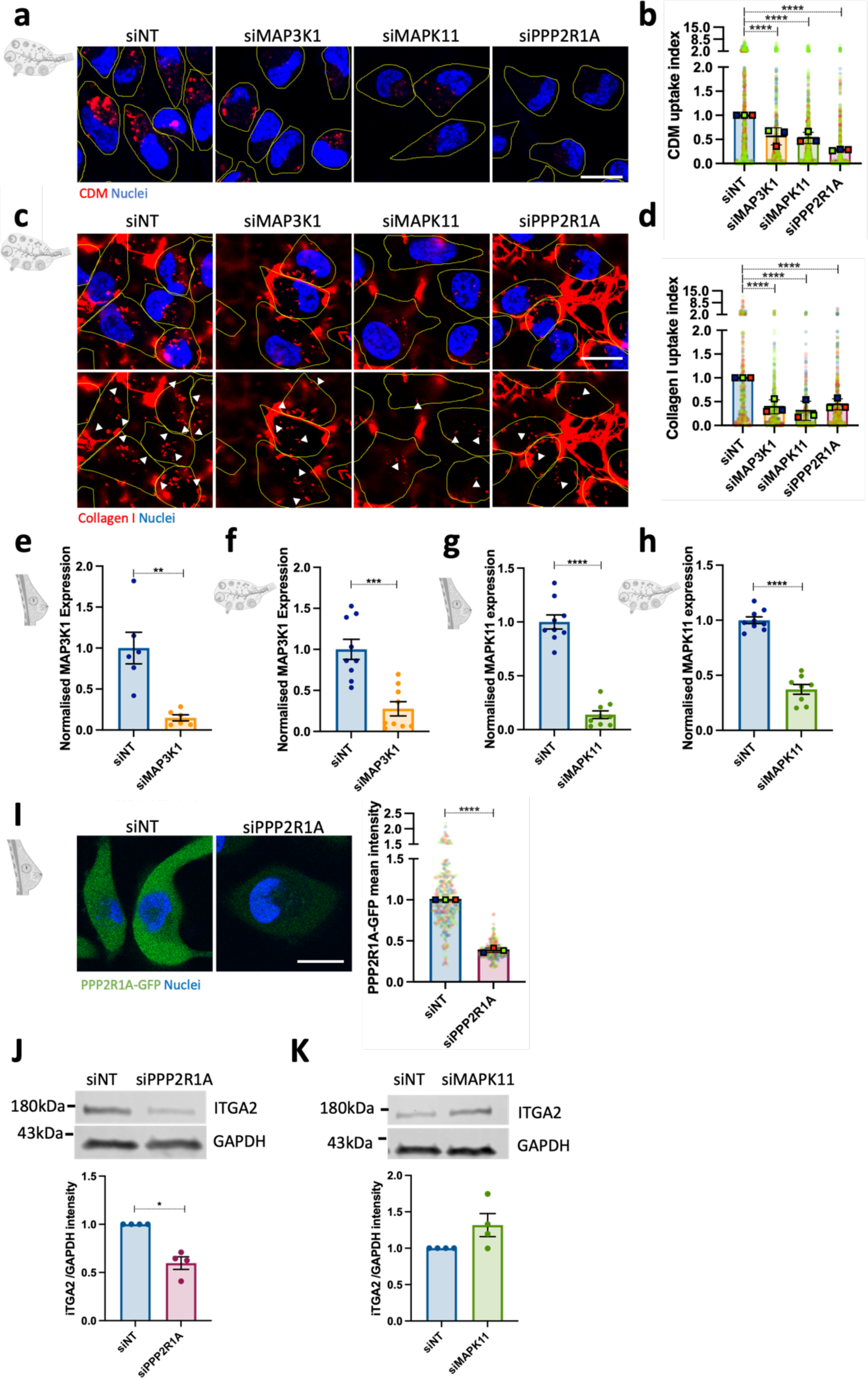
MAP3K1, MAPK11 and PPP2R1A knockdown decreased ECM internalisation. **a,** A2780-Rab25 cells were transfected with an siRNA targeting MAPK3K1 (siMAP3K1), an siRNA targeting MAPK11 (siMAPK11), an siRNA targeting PPP2R1A (siPPP2RlA) or a non-targeting siRNA control (siNT), seeded on pHrodo-labelled CDM for 6h, stained with 1μg/ml Hoechst and imaged live. Scale bar, 20μm. **b,** CDM uptake index was calculated with Image J. Values represented are normalised mean + SD from N=3 independent experiments; ****p<0.0001; Kruskal-Wallis test, **c,** A2780-Rab25 cells transfected as in a, seeded on 1mg/ml collagen I, labelled with NHS-Alexa fluor 555, for 6h, fixed and stained for nuclei. Scale bar, 20μm. **d,** Collagen I uptake index was calculated with Image J. Values represented are normalised mean + SD from N=3 independent experiments; ****p<0.0001; Kruskal-Wallis test. **e,f,** MDA-MB-231 (e) and A2780-Rab25 (f) cells were transfected with an siRNA targeting MAP3K1 (siMAP3K1) or a non-targeting siRNA control (siNT), RNA was extracted and MAP3K1 expression was quantified by qPCR. Normalised data from N=3 independent experiments; **p=0.0022; ***p=0.0003; Mann-Whitney test. **g,h,** MDA-MB-231 (g) and A2780-Rab25 (h) cells were transfected with an siRNA targeting MAPK11 (siMAPK11) or a non-targeting siRNA control (siNT), RNA was extracted and MAPK11 expression was quantified by qPCR. Normalised data from N=3 independent experiments; ****p<0.0001; Kruskal-Wallis test, i, MDA-MB-231 cells overexpressing PPP2R1A-GFP were transfected with an siRNA targeting PPP2R1A (siPPP2RlA) or a non-targeting siRNA control (siNT), fixed and stained for nuclei. GFP normalised mean intensity + SD from N=3 independent experiments is shown; ****p<0.0001; Mann-Whitney test. **J,k,** MDA-MB-231 cells were transfected with an siRNA targeting MAPK11 (siMAPK11, j), an siRNA targeting PPP2R1A (siPPP2 R1A, k) or a non-targeting siRNA control (siNT), α2 integrin (ITGA2) and GAPDH protein levels was quantified by Western Blotting. Data are presented as the normalised mean ± SD; N=4 independent experiments; *p=0.0286; Mann-Whitney test.

**Extended data Fig. 6.**
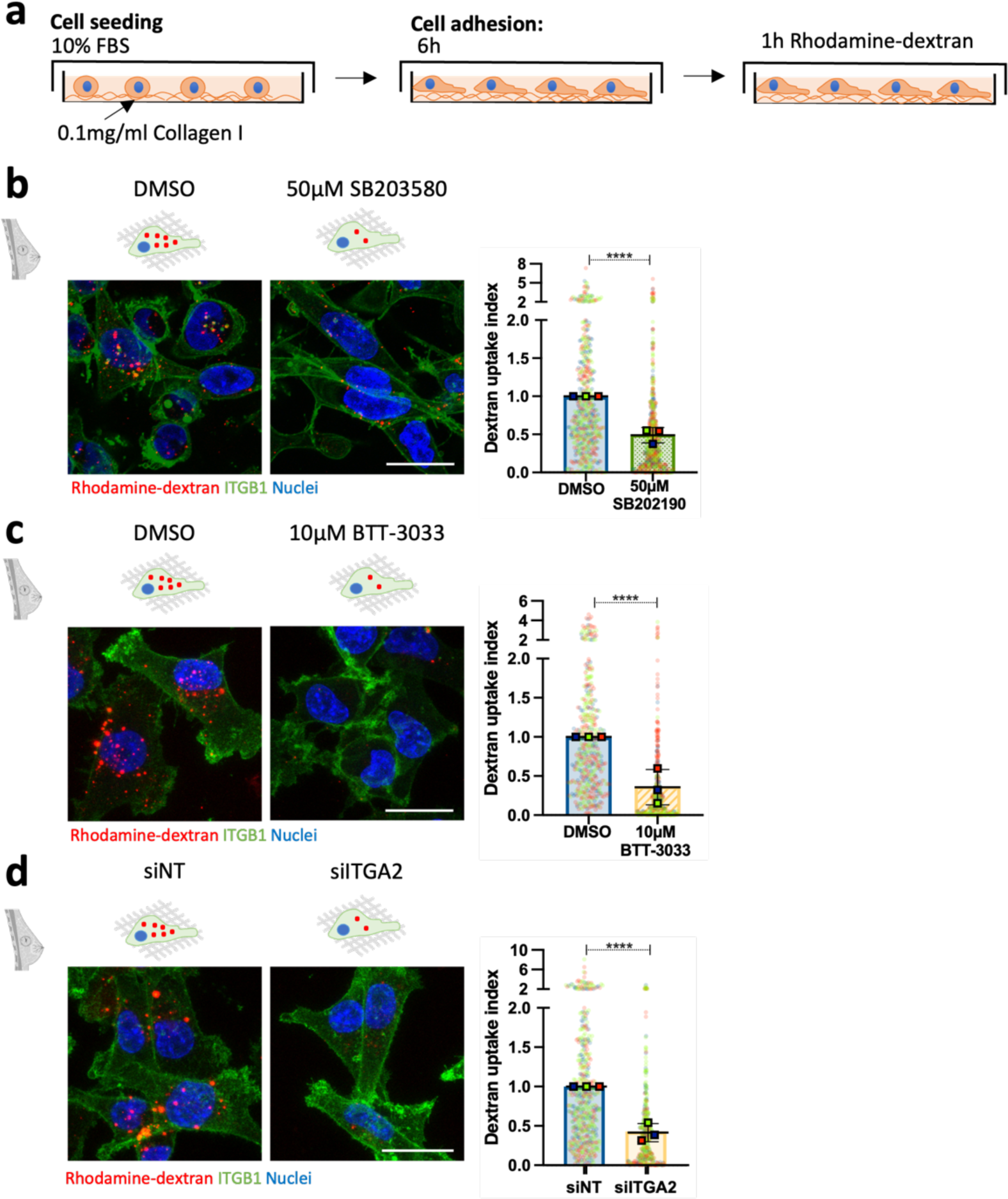
p38 and α2 integrin inhibition reduced macropinocytosis of soluble dextran. **a,** Schematic representation of the experimental set up. **b,** MDA-MB-231 cells were seeded on 0.1mg/ml collagen I for 6h, pre-treated with DMSO (vehicle) or S0µM SB202190 for 30 minutes, incubated with 0.2mg/ml rhodamine-dextran (red) for lh in the presence of DMSO or SB202190, fixed and stained for β1 integrin (ITGB1) and nuclei. Scale bar, 20µm. Dextran uptake index was measured with Image J. Data are presented as the normalised mean± SD; N=3 independent experiments. ****p<0.0001; Mann-Whitney test. **c,** MDA­ MB-231 cells were seeded on 0.1mg/ml collagen I for Gh in the presence of DMSO (vehicle) or 10μM BTT-3033, incubated with 0.2mg/ml rhodamine-dextran (red) for lh, fixed and stained for p1 integrin (ITGB1) and nuclei. Scale bar, 20µm. Dextran uptake index was measured with Image J. Data are presented as the normalised mean ± SD; N=3 independent experiments. ****p<0.0001; Mann-Whitney test. **d,** MDA-MB-231 cells were transfected with an siRNA targeting α2 integrin (silTGA2) or a non-targeting siRNA control (siNT), seeded on 0.1mg/ml collagen I for 6h, incubated with 0.2mg/ml Rhodamine-dextran (red) for 1hr, fixed and stained for β1 integrin (ITGB1) and nuclei. Scale bar, 20µm. Data are presented as the normalised mean± SD; N=3 independent experiments. ****p<0.0001; Mann-Whitney test.

**Extended data Fig. 7.**
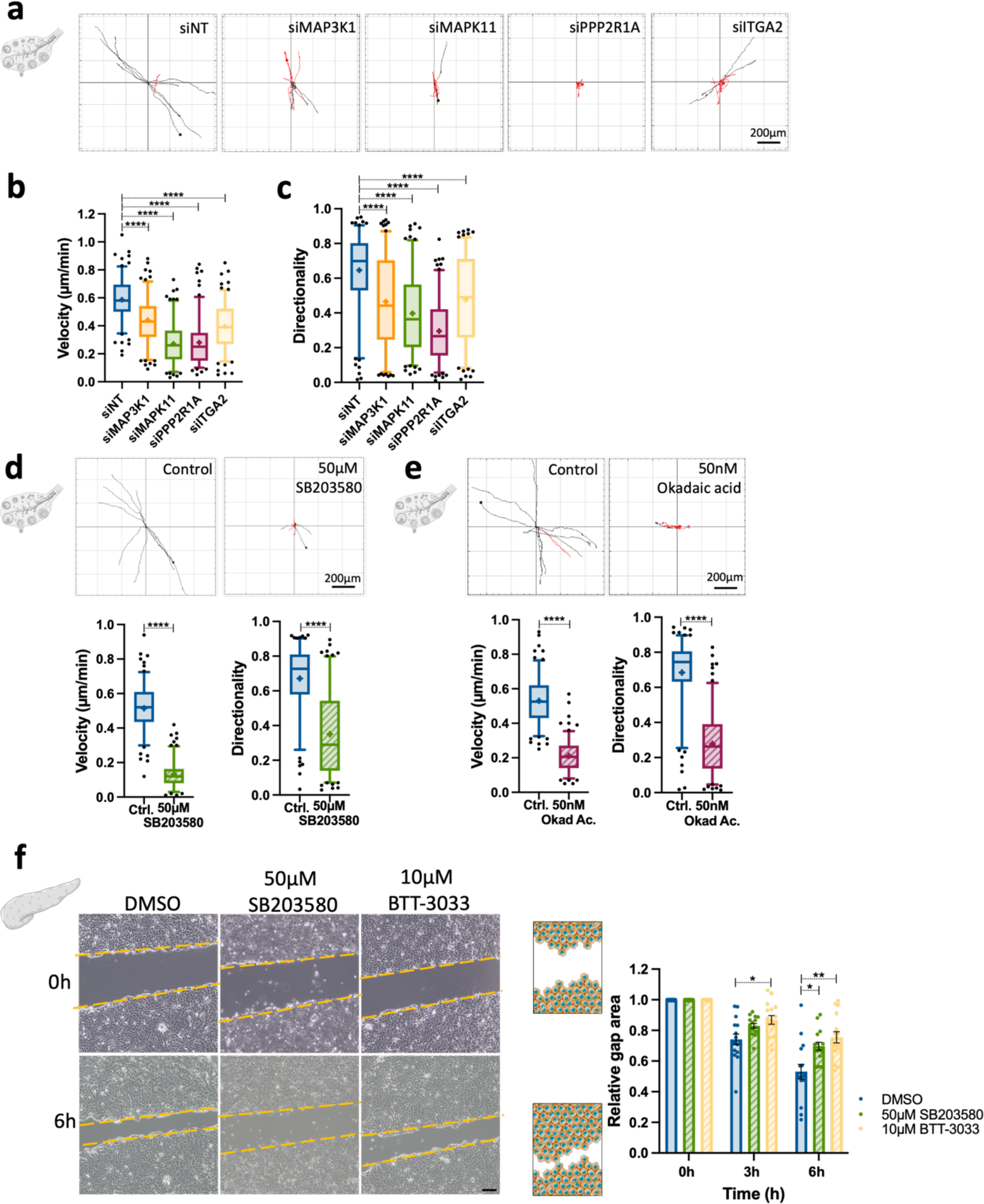
Regulators of ECM internalisation were required for ovarian and pancreatic cancer cell migration. **a-c,** A2780-Rab25 cells were transfected with an siRNA targeting MAP3K1 (siMAP3K1), an siRNA targeting MAPK11 (siMAPK11), an siRNA targeting PPP2R1A (siPPP2RlA), an siRNA targeting α2 integrin (si-ITGA2) or a non-targeting siRNA control (siNT), seeded on CDM for 6hr and imaged live with a 10X Nikon Inverted Ti eclipse with Oko-lab environmental control chamber for 17h. Spider plots show the migration paths of manually tracked cells (directionality >0.5 in black, <0.5 in red). Box and whisker plots represent 5-95 percentile, + represents the mean, dots are <5% and >95%; N= 3 independent experiments. ****p <0.0001; Kruskal Wallis test, **d,** A2780-Rab25 cells were seeded on CDM for 6hr in the presence of DMSO (Ctrl.) or 50μM SB203580 and imaged live with a 10X Nikon Inverted Ti eclipse with Oko-lab environmental control chamber for 17h. Spider plots show the migration paths of manually tracked cells (directionality >0.5 in black, <0.5 in red). Box and whisker plots represent 5-95 percentile, + represents the mean, dots are <5% and >95%; N= 3 independent experiments. ****p <0.0001; Kruskal Wallis test, **e,** A2780-Rab25 cells were seeded on CDM for 6hr in the presence of the vehicle (water, Ctrl.) and 50nM Okadaic acid (Okad Ac.) and imaged live with a 10X Nikon Inverted Ti eclipse with Oko-lab environmental control chamber for 17h. Spider plots show the migration paths of manually tracked cells (directionality >0.5 in black, <0.5 in red). Box and whisker plots represent 5-95 percentile, + represents the mean, dots are <5% and >95%; N= 3 independent experiments. ****p <0.0001; Kruskal Wallis test, **f,** SW1990 cell confluent monolayers were scratched and overlaid with 0.5mg/ml collagen I and cells were imaged at Oh, 3h and 6h. Yellow lines indicate the wound edges. The bar graph shows the normalised relative gap area ± SEM. N=4 independent experiments. *p<0.0213, **p=0.0025; 2way ANOVA.

**Extended data Fig. 8.**
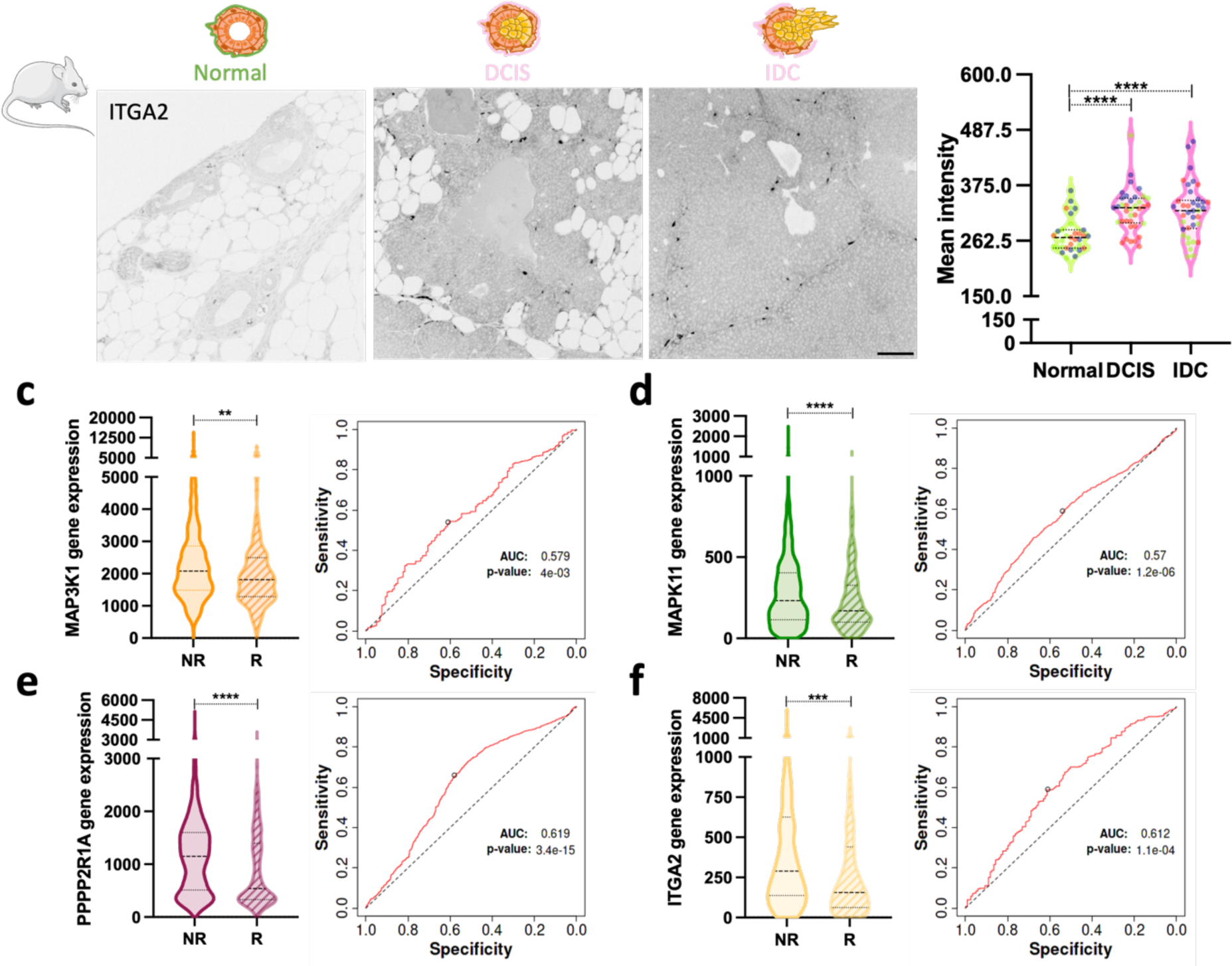
α2 integrin is upregulated in mouse mammary tumours and MAP3K1, MAPK11, PPP2R1A and α2 integrin expression is higher in chemotherapy-resistant breast cancer patients. **a,** Tissue sections from polyoma middle T-derived mouse mammary tumours were stained for α2 integrin (ITGA2). Scale bar, 50μm. **b,** α2 integrin mean intensity was quantified with Image J; N=3 independent experiments.****p <0.0001; Kruskal-Wallis test, **c-f,** RNA sequencing data and ROC analysis for MAP3K1 (a), MAPK11 (b), PPP2R1A (c) and α2 integrin (ITGA2, d) from chemoresistant (non-responder (NR)) and chemosensitive (responder (R)) breast cancer tumours.**p =0.0090; ***p =0.0002; ****p <0.0001; Mann-Whitney test.

**Supplementary video 1|MDA-MB-231 cells internalised and acidified collagen I.** MDA-MB-231 cells were seeded on NHS-fluorescein (green) and pHrodo-labelled 1mg/ml collagen I (magenta), cells were allowed to adhere for 30 minutes before being imaged live for 5h every minute with a 63X oil objective Zeiss LSM980 Airyscan 2 microscope with environmental chamber. Video corresponding to Extended data Fig. 1e.

**Supplementary video 2|MDA-MB-231 cells internalised collagen I during cell migration.** MDA-MB-231 cells were seeded on NHS-fluorescein (green) and pHrodo-labelled 1mg/ml collagen I (magenta), cells were allowed to adhere for 30 minutes and imaged live for 5h every minute with a 63X oil objective Zeiss LSM980 Airyscan 2 microscope with environmental chamber. Video corresponding to Fig. 5a.

**Supplementary video 3|MDA-MB-231 cells internalised CDM during cell migration.** MDA-MB-231 cells were seeded on pHrodo-labelled CDM (red) for 6h before imaging for an additional period of 6h with a 10X objective from a Nikon Inverted Ti eclipse with Oko-lab environmental control chamber. Video corresponding to Fig. 5b.

**Supplementary table 1|** Raw data values for Matrigel uptake index. Data were normalised between NT5+BafA1 (−1) and NT5 (0). Hits are shown in blue, while controls are in green.

**Supplementary table 2|** Reactome analysis of the positive and negative regulator hits obtained in the screening.

**Supplementary table 3|** Data were normalised between NT5+BafA1 (−1) and NT5 (0). The table shows kinome and phosphatome results and deconvolution of 4 individual siRNAs (2 biological replicates, 2 technical replicates per biological replicate). Validated positive regulators are highlighted in orange, while negative regulators are in green. Controls (ITGB1 and PAK1) are in dark red, NT5+BafA1 is in pink and NT5 is blue.

**Supplementary table 4|**Cell seeding values for generation of cell derived matrices.

**Supplementary table 5|**siRNA information.

## Notes

### Competing Interest Statement

The authors have declared no competing interest.

